# Hierarchical cortical plasticity in congenital sight impairment

**DOI:** 10.1101/2024.07.04.602138

**Authors:** Roni O. Maimon-Mor, Mahtab Farahbakhsh, Nicholas Hedger, Andrew T. Rider, Elaine J. Anderson, Geraint Rees, Tomas Knapen, Michel Michaelides, Tessa M. Dekker

## Abstract

A robust learning system balances adaptability to new experiences with stability of its foundational architecture. To investigate how the human brain implements this we combined structural imaging with population receptive field (pRF) and connective field (CF) modelling to study plasticity and stability across hierarchical processing stages in visual cortex. We compared the rod system of individuals born with rod-only photoreceptor inputs (achromatopsia) to the typically developed rod system, allowing us to dissociate impacts of life-long versus transient responses to altered input. Cortical input stages (V1) exhibited high stability, with input-deprived cortex showing no substantial retinotopic remapping and exhibiting structural hallmarks of deprivation. However, downstream cortex (V3) showed plasticity in how it read out these inputs. In achromats, more cortical resources in V3 were dedicated to sampling input from the near-fovea, and V3 neuronal populations aggregated information across smaller areas along the V1 cortical surface. We propose that these hierarchical dynamics reflect a broader principle of brain plasticity, in which later stages adapt to atypical input by altering how they sample earlier stages. This has important implications for emerging sight-rescue therapies, where restored input must be interpreted by a system that has already adjusted to its absence.

## Introduction

Neural systems need to be robust on the one hand but have flexibility to adapt to experience on the other. This places opposing pressures on the formation of the brain’s neural architecture. Historically, there have been two central contrasting views: early studies suggested that the brain exhibits remarkable plasticity, with the capacity for significant rewiring and reorganisation in response to new or altered experiences (Baker et al., 2005, 2008; Baseler et al., 2002; Calford et al., 2005; Dilks et al., 2009; Ferreira et al., 2016; Schumacher et al., 2008). More recent work portrays the brain as a fundamentally stable structure resistant to functional reorganisation even when primary sensory inputs are lost (Adams et al., 2007; Baseler et al., 2011; Goesaert et al., 2014; Molz et al., 2023; Ritter et al., 2019). Our study reframes this debate by investigating how adaptations of higher-order processing stages can compensate for stability of sensor-to-cortex connections, offering a new perspective on neuroplasticity through the lens of hierarchical network dynamics.

In the visual domain, the focal point of the debate on plasticity and stability has hinged on the extent to which retinal input deprivation can drive local reorganisation in early visual cortex, for example, for deprived tissue to take on inputs from spared retinal locations (Adams et al., 2007; Baker et al., 2005, 2008; Baseler et al., 2002, 2011; Calford et al., 2005; Dilks et al., 2009; Dumoulin & Knapen, 2018; Ferreira et al., 2016; Goesaert et al., 2014; Haak et al., 2015; Molz et al., 2023; Ritter et al., 2019; Schumacher et al., 2008). In reality visual impairment is a more global phenomenon, affecting all levels of visual processing, with complex dynamics beyond constricted local retinocortical projection zones (Carvalho et al., 2019; Masuda et al., 2021). Investigating these dynamics is crucial for understanding neural plasticity bottlenecks of regenerative treatments under trial in human patients (Fischer et al., 2020; Georgiou et al., 2021; Michaelides et al., 2023). In these treatments, signal is recovered at the level of the sensory organ, here the eye. If tissue at any downstream processing stage has deteriorated due to inactivation or has adopted new roles, it may be less responsive to its original signals. For ground-breaking sight recovery treatments to reach their full potential, understanding these cascading dynamics in the human brain is now paramount.

We revisit the debate on neuroplasticity and stability by investigating adaptations to altered sensory input across hierarchical cortical processing stages, using the inherited retinal disease achromatopsia as a model. Achromatopsia (also known as rod monochromacy) causes cone photoreceptors in the retina to be inactive from birth (Aboshiha et al., 2014). This means visual development is purely driven by rod photoreceptors. Rod-mediated vision causes low acuity, a blind spot (scotoma) in the rod-free retina of the fovea, nystagmus (involuntary eye-movements), and no colour vision. In most studies on how the brain adapts to sight loss, a major challenge is accurately matching the sensory input experienced by patients in control individuals with typical sight (Brown et al., 2021; Jones et al., 2020; Macnamara et al., 2021). This matching is critical for distinguishing neural adaptations to long-term altered visual experience from normal transient responses to varying visual input (Binda et al., 2013; Carvalho et al., 2021; Haak et al., 2012; Hummer et al., 2018). While normally sighted individuals rely on cone vision most of their waking time, rod only vision can also naturally occur or be induced experimentally (e.g. Barton & Brewer, 2015). This makes it possible to directly compare lifelong rod-only vision in achromatopsia with transient rod-only experiences in normal sight, distinguishing the long-term neural impacts of altered sensory input from temporary changes within the typical system’s range of flexibility.

It remains unclear how the brain adapts to altered retinal input in achromatopsia. Initially, a functional MRI study showed that areas of the early visual cortex, typically receiving only foveal cone input, might reorganise to process input from peripheral rod-containing regions (Baseler et al., 2002). This suggests potential cortical plasticity. However, recent findings by Molz et al. (2023) present a conflicting view, showing no reorganisation in the deprived primary visual cortex of achromats, and structural signs of deprivation (Molz et al., 2022). This suggests functional stability. Here, we reasoned that both stability and plasticity in a deprived visual system can co-exist across different cortical processing stages; A high degree of stability in sensor-cortex connections (Haak et al., 2015; Molz et al., 2023) may be offset by plasticity in downstream regions in the hierarchy, which may exhibit more scope to flexibly adapt and optimise the use of available input.

To test this prediction, we drew on a uniquely large sample of achromats and controls. We first characterised spatial tuning and remapping of V1 in a cone-input deprived visual system to resolve ambiguity of earlier findings. We then used a connective field (CF) modelling approach to assess sampling of V1 information by higher order visual cortex (V3). To truly distinguish long-term adaptations from transient changes to altered input, we took a different approach than prior studies. Rather than only manipulating luminance levels, we utilised the method of silent-substitution (Estévez & Spekreijse, 1982) to create photoreceptor-specific population receptive field (pRF) mapping that only stimulates rods. This was essential because pRF estimates are stimulus-dependent: they reflect the aggregate response of neuronal populations within a voxel and can vary both with the retinal input generated by a stimulus and with the subsets of neurons contributing to the measured signal. By obtaining well-matched, high-quality rod-driven maps from both achromats and controls, we were able to make an unbiased comparison of rod-mediated responses across groups.

Using these specialised stimuli to characterise the impacts of purely rod-based vision on the visual systems of achromats and normally sighted controls, we first show that V1 retinotopy presents the hallmarks of a highly robust system. There is no evident remapping of cortical regions that are input-deprived under rod-only viewing circumstances, despite their life-long altered input in achromatopsia. Some changes in pRF tuning (increased pRF size) outside the deprived region might be present. However, with our current data it is difficult to identify whether the origin of this difference comes from true tuning changes or measurement confounds. We found that the robustness to long-term altered input in V1, is paired with reorganisation of retinotopic read-out by higher-order visual cortex (V3). A comparison of the visual space organization of V1’s connectivity with V3, reveals a shift in the distribution of resources in achromats: more cortical resources in V3 are dedicated to sampling the V1 cortical surface that encodes 1°-4° visual angles - the most foveal areas accessible to rods. Vertices in V3 also sample from fewer V1 vertices, i.e., show a reduction in CF size, resulting in more normalised receptive-field extent in V3. These changes in the rod-based visual system of achromats could act to compensate for the absence of two features of the cone system that the rod system lacks: its high-density foveal sampling and its overall higher resolution. This may reflect an overarching broader organising principle where hierarchical optimisation processes allow the brain to make more efficient use of the available (in this case atypical) input.

## Results

In three results sections, we characterise the interplay between neural plasticity and stability in achromatopsia across three complementary neural markers: brain structure, spatial tuning characteristics of the functional MRI signal in V1, and the hierarchical retinotopic connectivity patterns that indicate how higher-order visual cortex (V3) samples information from V1.

### Structural analysis

As a proxy of structural integrity, we compared cortical thickness across 8 eccentricity segments of V1 (0°-1°,1°-2°,2°-3°, etc.; see Figure 1B) in a large sample of 31 achromats and 100 controls aged 6-35 years. Data were collated over two scanners. We expected changes in achromats to be largest in the segments representing the central 0°-2° eccentricities of the visual field, encompassing the retinal ‘rod-free’ zone, an oval of approximately 1.25°x0.96° in size (Curcio et al., 1990). V1 segments were defined using the Benson atlas (Benson & Winawer, 2018). The atlas fit procedure uses cortical landmarks on individual brains to predict the viewing eccentricity to which each cortical vertex is tuned. This allows us to identify where the representation of these eccentricities would have been given typical development, and investigate what they represent in atypical development.

**Figure 1.**
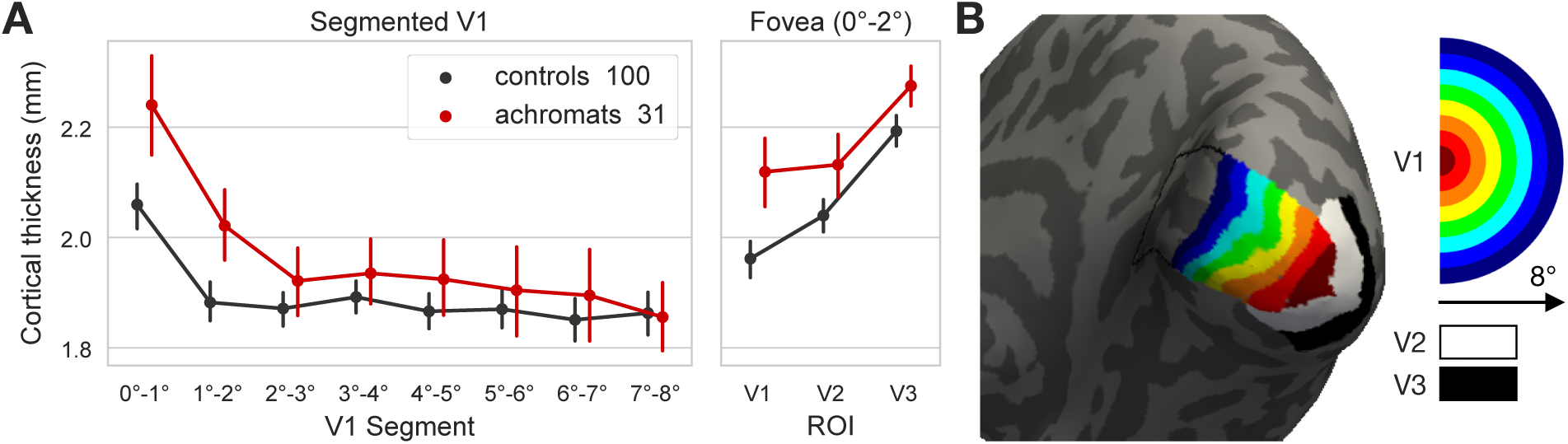
Structural properties of V1 in achromats and controls. (**A**) Mean V1 cortical thickness of each eccentricity based V1 segment in millimetres (mm). Cortical thickness was increased in achromats (n=31) compared to controls (n=100) only for foveal projection segments 0°-2° and not in more eccentric segments 2°-8°. This pattern was trending in V2 and significant in V3. Error bars are 95% confidence intervals (**B**) Visualisation of regions of interest in one control participant. Segmentation of V1 to 1° eccentricity bins, and the foveal projection zone (0°-2°) of V2 and V3 was based on the Benson retinotopy atlas (Benson and Winawer, 2018). The half circle on the right shows, in visual space, the area represented by each segment.

Figure 1A shows an increase in cortical thickness in achromats compared to controls only in the foveal projection zone (Controlling for scanner, overall cortical thickness, and age; 0°-1° ANCOVA group effect: *F*_(1,126)=_9.05, *p_unc_*=0.003; 1°-2°: *F*_(1,126)=_7.79, *p_unc_*=0.006). This increase was not present in more eccentric parts of V1 (*p*_unc_>0.2 for all). A further comparison of the cortical thickness in the foveal projection zones (0°-2°) in V2 and V3 (Figure 1A right), revealed a trend toward significance in V2 (*F*_(1,126)_=3.07, *p*=0.08) and a significant effect in V3 (*F*_(1,126)_=6.32, *p*=0.01). As thicker cortex in early visual areas is a hallmark of visual deprivation from birth (Jiang et al., 2009; Park et al., 2009), this indicates that the foveal projection zones in achromats may lack retinal input. This provides evidence for stability, because if these cortical areas had substantially reorganised to receive more peripheral retinal input, we would not expect to see this marker of deprivation.

### pRF properties of V1

We next test whether the functional visual-spatial tuning of V1 in a system that has developed using only rod input (i.e., achromats) remains stable or adapts, using population receptive field modelling. Our analyses focus on two types of plasticity. First, we ask what happens to the V1 cortex normally receiving only cone-mediated input from the foveal ‘rod-free’ zone, which shows structural abnormalities. Second, we test for changes in rod-selective visuospatial tuning across V1 (beyond the 0°-2° ‘rod-free’ fovea), in cortex which normally receives combinations of rod and cone input but develops with only rod inputs in achromats.

V1 receives retinotopically organised input, with adjacent neurons in the cortex receiving signals from adjacent neurons in the retina. Population receptive field modelling links retinotopic voxels in visual cortex to the locations in the visual field they represent (Dumoulin & Wandell, 2008). This is achieved by fitting population receptive fields estimates (pRFs; i.e., Gaussian tuning functions) to BOLD responses evoked by stimuli systematically traversing the visual field. To compare rod-mediated cortex function in achromatopsia to normally developed rod-responses under matched viewing circumstances, we used the silent substitution method (Estévez & Spekreijse, 1982) and low luminance presentation which allowed us to generate stimuli with a rod-isolating contrast (“rod-selective condition”). We also included another stimulus condition that, in a typical retina, stimulates cones and rods equally (“non-selective condition”). This allowed us to assess how rod-mediated measures in both groups compare to more typical pRF measures driven by cones in normally sighted, and whether differences across achromats and controls remain consistent across different visual stimulus properties.

### Coverage and pRF properties of the foveal projection zone

We first compared retinotopic tuning in cortex containing the rod-free foveal projection zone (atlas defined 0°-2°) across controls and achromats. In normally developed retinotopic rod maps, the cortical projection of the foveal rod-free zone, appears as a central “hole” (Figure 2B; Barton & Brewer, 2015). To test if this normally silent area fills in with reorganised retinotopic visual activity in achromats as has previously been reported (Baseler et al., 2002), we compared the percentage of cortex that contains reliable pRFs (coverage; meaning where pRFs could reliably be estimated from visually-driven signal). This is shown in Figure 2 for pRF model fits of R^2^ > 0.1, but effects were similar at R^2^ > 0.2 and 0.3 (see Supplement 6).

**Figure 2.**
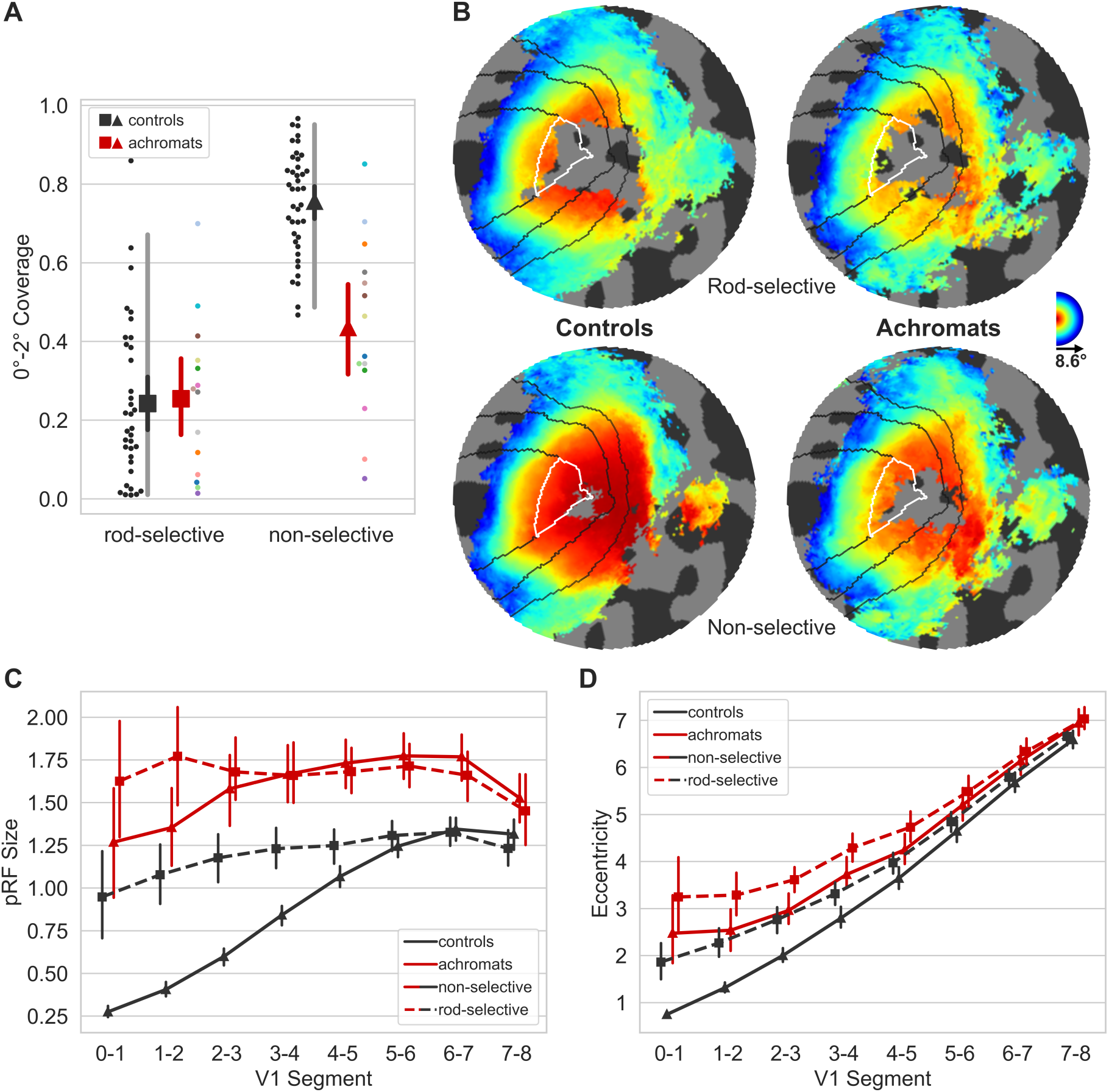
pRF properties of V1 in achromats and controls. (**A**) Foveal projection zone (0°-2°) coverage. Coverage is the fraction of vertices within the foveal confluence that contain visual information (i.e., pRF R^2^>0.1). Achromats (red markers) are not significantly different from controls (black markers) in the rod-selective condition (square markers), and do not show complete filling in even in the non-selective condition with higher contrast (triangle markers). Triangle markers: group mean for the non-selective condition, square markers: group mean for the rod-selective condition. Black markers: controls group mean, red markers: achromat group means. Black/red vertical lines around the mean: 95% confidence interval, grey shaded lines: 2.5%-97.5% percentile interval (only in controls). For achromats, participants’ dots are colour-coded to match across conditions and measures (**B**) Group eccentricity maps plotted on a flat surface for each condition (in standard fsaverage space), presenting the group median value of each vertex. For visualisation purposes, vertices that were absent in more than 30% of the group were left blank in each condition. Black lines denote atlas-based delineations of V1,2,3; white lines mark the foveal projection zone (0°-2°) in V1. As shown in (A) we see no evidence of ‘filling in’ of the rod scotoma in achromats. (**C**) pRF sizes along the atlas V1 segments. Achromats show a consistent increase in pRF size compared to controls. (**D**) Eccentricity of modelled pRFs along the atlas V1 segments. Achromats show an increase in eccentricity compared to controls, however that shift is not consistent across the two stimulus types. Vertical lines in C&D represent 95% confidence intervals.

In the rod-selective condition, the coverage in the 0°-2° region of interest was near-identical across achromats and controls (on average 24% in controls, and 25% in achromats, *t_(47)_ = 0.19, p=0.85, BF_10_=0.31*; see Figure 2A&B), suggesting no substantial filling in of the foveal projection zone despite life-long rod-only vision. For non-selective stimuli targeting both cones and rods, controls exhibit greater visual coverage (75% on average) compared to achromats, who are missing functional cones (43% on average) (Mann *U_(52)_=63, p<0.0001; unequal variance reverted to non-parametric*). Achromats show enhanced coverage under non-selective compared to rod-selective conditions, likely due to increased luminance yielding stronger activations. We note that two achromats show potential filling in, with a foveal activation coverage beyond the 95% predictive interval of the normally sighted rod gap. Their retinotopic profiles (see Supplement 1) suggest this may reflect individual differences in retinal foveal photoreceptor distribution profiles rather than cortical remapping. This highlights the importance of characterising variability in atypically developing populations through larger samples rather than case studies.

Crucially, data quality was matched across groups, with no differences in pRF model fits. Both prior to any R^2^ filtering (*t*_(48)_=0.53, *p*=0.60, BF=0.34; see Figure S2b), or after filtering R^2^ > 0.1 (*t*_(48)_=−0.06, *p*=0.95, BF=0.31). It is therefore unlikely that any coverage inside the rod “hole” in achromats was masked by low data-quality differences between the groups. In addition, it is unlikely that these findings can be explained by pRF fitting-related artefacts (Binda et al. 2013), as modelling the rod-scotoma in the pRF fitting procedure led to similar results (see methods and Supplement 3).

Another potential confound in our findings is fixation instability. In pRF mapping, which is usually conducted under photopic (cone-dominant) conditions, unstable fixation can cause a signal drop in the foveal projection zone. As expected due to nystagmus, the achromatopsia group showed higher fixation instability compared to controls (*rod-selective: t_(9.08)_=−3.19, p=0.01*; *non-selective*: *t_(9.41)_=−4.88, p<0.001*; *degrees-of-freedom corrected for unequal-variance*; see Supplement Figure S2a). However, several lines of evidence suggest this instability cannot fully account for the lack of “filling in” in achromats. First, within the achromat group, we found no correlation between fixation stability and coverage (*rod-selective: spearman-r_(8)_ = −0.36, p=0.31; non-selective spearman-r_(8)_=0.07,p=0.85)*; Individuals with more stable, control-like fixation did not show more signal inside the scotoma (see Supplement 2). Second, in adults with achromatopsia, typically with less severe nystagmus (Kohl et al., 1993), two recent studies also found absence of filling in (Anderson et al., 2024; Molz et al., 2023).

So, while we cannot fully exclude nystagmus masking foveal signals in the cortex of some patients, this converging evidence from structural and functional MRI measures across different studies and groups, strongly suggests that the deprived cortex does not substantially ‘fill in’ with peripheral rod inputs in achromatopsia.

Together these results provide strong evidence for stability in the face of life-long deprivation, i.e., the conservation of the sensory topography.

### Altered rod representation in V1 - pRF size and eccentricity

We next tested for differences in spatial tuning across V1 regions that typically receive both rod and cone inputs but develop with only rod inputs in achromatopsia. Figure 2C shows that pRF sizes in achromats are systematically larger compared to controls across all eccentricities. This holds true for both the non-selective (V1 segments: 2°-3°: *t*_(52)_=*12.57, p_unc_<0.001*; 3°-4°: *t*_(52)_*=11.12, p_unc_<0.001*; 4°-5°: *t*_(52)_=*9.54, p_unc_<0.001*; 5°-6°: *t*_(52)_=*7.52, p_unc_<0.001*; 6°-7°: *t*_(52)_*=5.76, p_unc_<0.001*; 7°-8°: *t*_(52)_*=2.36, p_unc_=0.022;* all segments remain significant after FDR correction), and rod-selective stimulus conditions (V1 segments: 2°-3°: *t_(48)_=3.83, p_unc_<0.001;* 3°-4°: *t_(48)_=3.67, p_unc_ <0.001*; 4°-5°: *t_(48)_=4.68, p_unc_ <0.001*; 5°-6°: *t_(48)_=4.74, p_unc_ <0.001*; 6°-7°: *t_(48)_=3.98, p_unc_ <0.001*; 7°-8°: *t_(48)_=2.00, p_unc_ =0.051*; all segments but the 7°-8° remain significant after FDR correction). In the non-selective condition, the smaller pRFs in controls are in line with the higher spatial resolution of the cone system, which is not active in the achromat group. These differences should therefore not be interpreted as direct evidence for altered neural tuning in achromatopsia. They instead reflect different retinal input to V1 and, likely, different neuronal populations in V1 engaged by cone-driven responses in controls versus rod-driven responses in achromats. However, for the rod-selective condition where both groups activate the same class of photoreceptors, the enlarged pRFs in achromats might indicate altered rod-based visual processing.

Larger pRFs indicate that neuronal populations in achromats’ V1 cortex, combine information across larger areas in visual space than in typically sighted controls. This could reflect true neural tuning differences as well as be driven by larger eye movement. However, fixation instability in achromats does not significantly correlate with pRF size in our sample (*rod-selective: spearman-r_(8)_ = −0.41, p=0.24; non-selective spearman-r_(8)_=−0.37,p=0.29*).

Figure 2D shows that pRF estimates in achromats have higher eccentricities across V1 compared to controls, spanning all tested V1 segments (group comparisons of the rod-selective conditions: 2°-3°: *t*_(48)_*=3.32, p_unc_=0.002;* 3°-4°: *t_(48)_=4.42, p_unc_<0.001;* 4°-5°: *t_(48)_=3.59, p_unc_=0.001*; 5°-6°: *t_(48)_=3.17, p_unc_=0.003*; 6°-7°: *t_(48)_=3.15, p_unc_=0.003*; 7°-8°: *t_(48)_=2.23, p_unc_=0.030*; all segments but the 7°-8° remain significant after FDR correction). This slight outward eccentricity shift could reflect true changes in neural tuning. However, eccentricity estimates also varied across stimulus conditions. In controls, such differences may be explained by the different photoreceptor populations driving the neural activity across the two conditions. In achromats, however, both conditions stimulated only rods using different luminance and contrast levels, yet the non-selective condition produced lower eccentricity estimates than the rod-selective condition. This suggests that the pRF position estimates are stimulus-dependent to some degree, even when the same photoreceptor class is stimulated. This difference was not explained by selective drop-out of vertices with smaller or more central pRFs, as the pattern remained identical when the analysis was restricted to vertices with significant pRF estimates in both stimulus conditions. Interestingly, the eccentricity estimates of the non-selective condition in achromats appear similar to those of the rod-selective condition in controls. This suggests that eccentricity estimates in the normal rod range can be obtained in achromats.

It has been shown that fitting artefacts around scotoma edges, can give rise to similar outward eccentricity shifts (Binda et al., 2013). However, when accounting for fitting artefacts around the foveal scotoma edge by modelling the rod-free zone during pRF fitting, pRF size and eccentricity differences remain unchanged (see Supplement 3). Finally, we found no significant correlations between gaze stability and the eccentricity shift (*rod-selective: spearman-r_(8)_ = 0.58, p=0.08; non-selective spearman-r_(8)_=0.09,p=0.8,* see Supplement 2).

Together, these analyses reveal subtle differences in how V1 of achromats responds to rod signals outside the foveal zone, which are consistent with results from other studies (Molz et al. 2023, Anderson et al. 2024). While we found no direct evidence that these are being driven by confounding factors such as eye-movements or fitting artefacts, more work is needed to understand the underlying processes that give rise to these shifts.

### Connective field modelling – hierarchical sampling

We next used connective field modelling to test whether V3’s sampling along the V1 cortical surface in achromatopsia remains stable given the evidence that the foveal part of V1 remains visually silent from birth. We test for two hypothetical types of hierarchical plasticity: shifts in where along the V1 surface V3 samples its information from (CF centre), and shifts in the extent of the area along the V1 surface that V3 samples from (CF size).

Higher-order visual regions receive retinotopic information from V1, with individual neurons sampling information across multiple neighbouring V1 neurons. Connective field (CF) modelling is a functional connectivity-based extension of pRF modelling which characterises this sampling process, based on time-series correlations. While pRF modelling is ‘stimulus referred’, estimating a voxel’s spatial tuning properties based on its response to stimulus variations, connective field modelling is ‘neural-referred’, estimating a voxel’s ‘connective field’ based on its correlations with aggregated voxels along the V1 surface. Responses throughout cortex are modelled as deriving from Gaussian shaped sampling profiles over the V1 cortical surface (connective fields), defined by a position (centre of the Gaussian) and size (Gaussian spread in mm (σ)). CF modelling can thus reveal the topographic profile of a brain region’s neural connectivity with V1, including the spatial location and extent of sampling. After estimating connective fields, the CF positions can be combined with the retinotopic map of V1 to ‘project’ the retinotopy of V1 onto the rest of the brain, thereby performing connectivity-derived retinotopic mapping. To create V3 connective-field-based eccentricity maps (Figure 3B), each vertex was assigned the Benson atlas eccentricity of the V1 segment of its connective field centre. CF modelling is especially informative in cases of visual deprivation. Time-series correlations can reflect not only visually evoked responses but also intrinsic fluctuations that flow across connected neural populations. CF modelling may therefore reveal preserved connectivity with regions such as the rod-free foveal projection zone, where little or no visual input is expected. Moreover, because CF modelling makes no assumptions about visual input, it is more robust than pRF modelling to confounds such as poor fixation and low visual responsiveness (Tangtartharakul et al., 2023). Here we focus on connective fields between V3 and V1. We omit V2, which shares a border with V1, to avoid inaccuracies in CF estimation due to mis-delineation.

**Figure 3.**
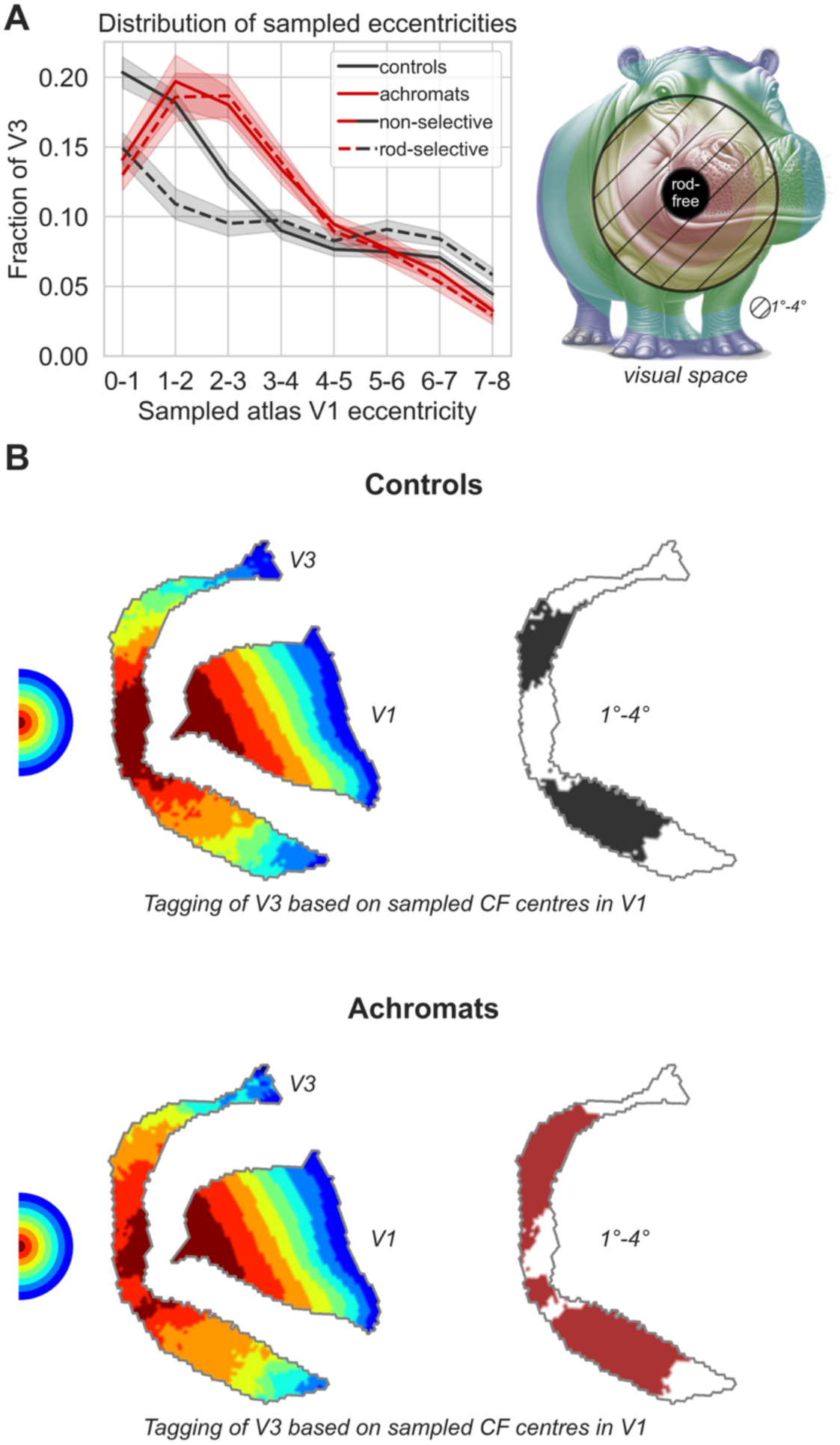
Connective field modelling (V3) in achromats and controls – sampled eccentricities. (**A**) Left: The fraction of all vertices within V3 that sample from V1, binned by sampled V1 atlas eccentricity. In achromats there is a shift towards greater sampling from the most foveal area that contains rods (1°-4°) of V1 relative to the normally sighted rod and cone systems. Shaded areas: standard errors of the mean (s.e.m). All V1 eccentricities are derived from the Benson Atlas. Right: A schematic illustration of the sampled eccentricities in visual space. Dashed area represents the area where group differences in sampling were found, 1°-4°. (**B**) Visualisation of V3 sampled eccentricities of V1 in the rod-selective condition. From left to right: Half circle represents the visual field, where each colour represents 1 visual degree. V1 is colour-coded into segments based on the atlas-assigned eccentricities. Each vertex in V3 is then coloured based on the centre of Gaussian connective field within the V1 surface. Values in V3 are based on group medians. To further highlight the group differences in the amount of cortical surface devoted to the 1°-4° V1 segments we visualise V3 again on the right, colouring in all vertices with a CF centre at the 1°-4° segments.

#### Shifts in V3 resource distribution along the V1 eccentricity map

When both rods and cones are stimulated in the normally sighted system, the largest proportion of V3 voxels is dedicated to sampling information from the V1 foveal projection (connective field centres in the 0°-1° degree range), with a gradual decrease in resources allocated to larger eccentricities (Figure 3A). When only rods are stimulated, foveal sampling is attenuated but the overall pattern of a gradual decrease in sampling from small to larger eccentricities remains. In achromats, the way in which V3 samples V1 is drastically different, with most V3 resources dedicated to sampling eccentricities at 1°-4° for both stimulation conditions (Figure 3A&B). When comparing the rod-selective condition in achromats to that of controls, we observe a shift towards greater V3 sampling from the V1 cortex encoding the (1°-4°) paracentral visual field (0°-1°: *t_(47)_=1.18, p_unc_=0.242*; 1°-2°: *t_(47)_=*4.03, *p_unc_*<0.001; 2°-3°: *t_(47)_=5.32, p_unc_<0.001*; 3°-4°: *t_(47)_=3.70, p_unc_=0.001*; 4°-5°: *t_(47)_=2.32, p_unc_=0.025*; 5°-6°: *t_(47)_=0.84, p_unc_=0.405*; 6°-7°: *t_(47)_=−0.97, p_unc_=0.337*; 7°-8°: *t_(47)_=1.93, p_unc_=0.060;* segments including 1°-4° remain significant after FDR correction). It is unlikely that these differences result from differences in measurement quality, because the goodness of fit (CF R^2^) was comparable across the two groups (Supplement 4). We therefore hypothesise that this shift in sampling reflects increased neural resources dedicated to the most fovea-adjacent information that is available to the rod-only system. Surprisingly, even though we found no evidence of visually-driven signal in the region on V1’s surface that normally takes foveal input (0°-1°) in achromats (see Figure 2), there is still a substantial portion of V3 vertices (15%) that sample from this region. This remaining foveal connectivity between V1 and V3 shows that there is also a degree of stability across retinotopic hierarchical sampling.

### Shifts in the V3 read-out aperture (CF size) from V1

Figure 4A shows that in normally sighted individuals, connective field size in V3 increases with eccentricity, whether stimulating both rods and cones at the same time, or only rods. This replicates findings by Haak et al., (2013) and Knapen (2021). In achromatopsia, however, V3 connective field sizes remain consistently small across all eccentricities, and are significantly smaller than in the normally sighted rod-system between 4°-6° eccentricity (comparing achromats vs control rod-selective condition: 0°-1°: *t_(44)_=1.07, p_unc_=0.289*; 1°-2°: *t_(45)_=−0.41, p_unc_=0.684*; *2*°*-3*°*: t_(45)_=−0.58, p_unc_=0.568*; 3°-4°: *t_(46)_=−1.70, p_unc_=0.097*; 4°-5°*: t_(47)_=−3.55, p_unc_=0.001*; 5°-6°: *t_(47)_=−2.63, p_unc_=0.011*; 6°-7°: *t_(47)_=−0.81, p_unc_=0.420*; 7°-8°: *t_(45)_=−0.73, p_unc_=0.469*; degrees of freedom vary as not all participants have vertices sampling all V1 segments). These smaller connective field sizes in V3 mean that V3 vertices pool across a smaller number of vertices along the V1 cortical surface. Visually driven signal could pull CF positions towards V1 regions that carry visual input, such as the scotoma border, causing a model-fitting bias around a scotoma edge. However, controls also lack visually driven input to the rod-free zone in the rod-selective condition. Any such bias should therefore affect both groups equally and cannot by itself explain the shift in V3 sampling we observe in achromats. Moreover, V3 vertices sampling from the immediate vicinity of the scotoma (1°-3°) show CF sizes comparable to controls. The significant reduction in CF size occurs only further in the periphery (4°-6°), in regions that are primarily stimulus-driven.

**Figure 4.**
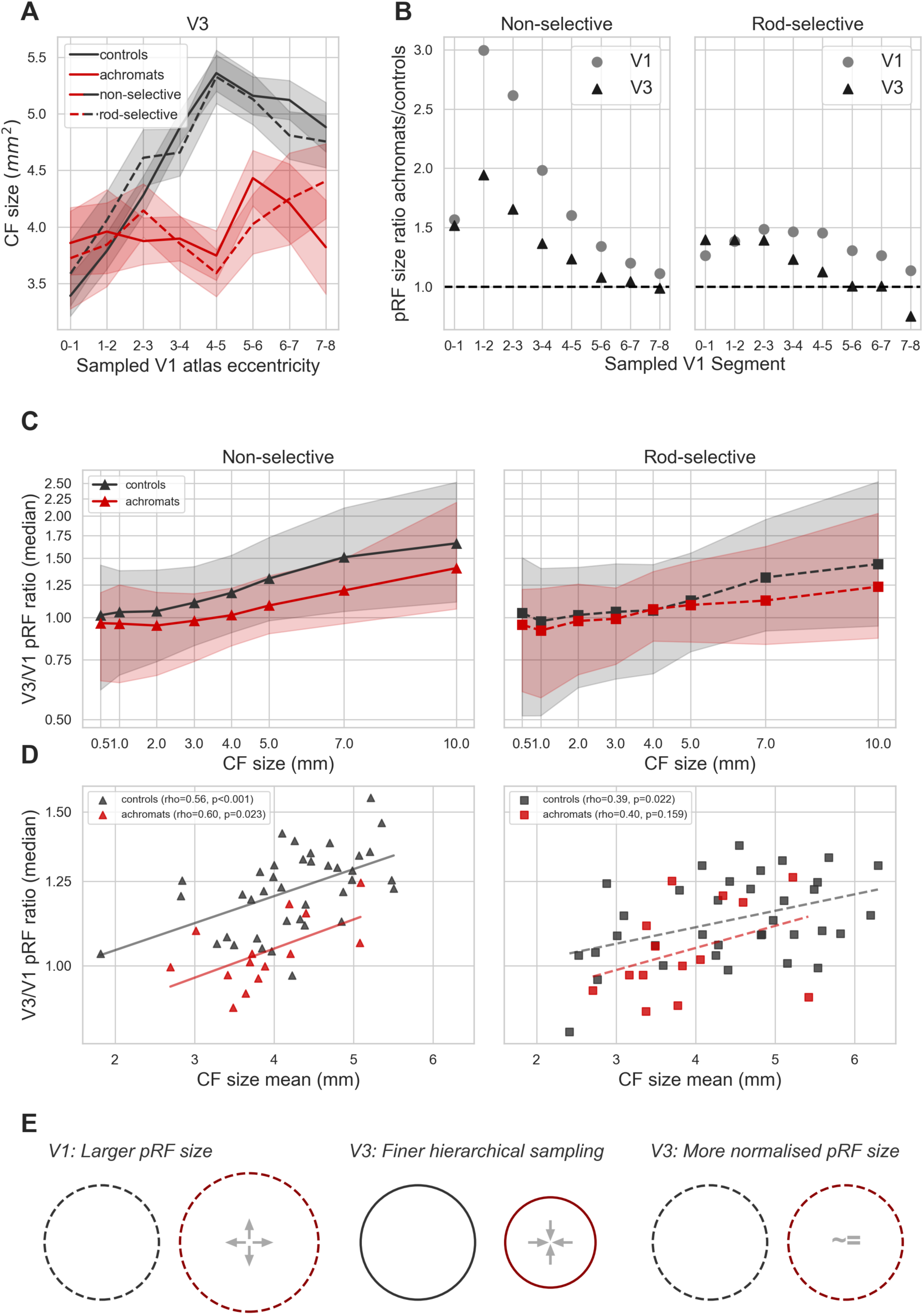
Connective field modelling (V3) in achromats and controls – connective field size and relationships with pRF size. (**A**) CF sizes in V3 in achromats and controls across the rod-selective and non-selective condition. Controls show an expected increase in CF size with sampled eccentricity while achromats show a flat pattern, with smaller CFs between 4°-6°. In both groups results are consistent across the two stimulation conditions. (**B**) Ratio of pRF size between achromats and controls in V1 (circles) and V3 (triangles). A ratio of 1 indicates the mean pRF size of achromats is equivalent to that of controls. Across both non-selective (left) and rod-selective (right) stimulus conditions, the ratio of pRF sizes in V3 is closer to 1 than in V1, pointing towards a possible normalising effect of CF size on pRF size. (**C**) Median V3/V1 pRF-size ratio (the ratio of a V3 vertex’s own pRF size to the pRF size of its CF centre in V1) as a function of CF size, for controls (black) and achromats (red), in the non-selective (left, triangles, solid line) and rod-selective (right, squares, dashed line) conditions. Shaded bands show the interquartile range. The ratio increases with CF size in both groups and conditions. (**D**) Subject-level relationship between pRF V3/V1 ratio and mean CF size. Plotted and correlated (spearman) in log-ratio space. (**E**) A summary of the proposed model of how smaller CF sizes in V3 of achromats might operate as a compensatory mechanism for increased pRF size in V1, resulting in normalized pRF size at the level of V3.

We next asked how this finer cortical sampling in V3 (smaller connective fields) aligns with visual processing as captured by pRF modelling. In V1, pRF sizes in achromats were significantly larger than in controls for both stimulus conditions, indicating coarser spatial tuning at the cortical input stage (Figure 2C). Finer sampling downstream cannot recover spatial detail that is already absent from the V1 representation. It can, however, constrain the spatial extent of downstream tuning: if V3 sampled the same extent of the V1 surface as in controls, the enlarged V1 pRFs would yield correspondingly enlarged V3 pRFs. By selectively sampling from a smaller area of the V1 surface (smaller CFs), V3 may reach a comparable extent of visual space from fewer V1 vertices, resulting in pRF sizes that are closer to control values than those observed in V1. If so, this process should result in a relative normalisation of pRF size in V3 compared to V1 (Figure 4E).

To test this prediction, we plotted the ratio of pRF sizes between achromats and controls, where a value of 1 indicates parity between the groups (Figure 4B). Consistent with the smaller CF sizes found in V3, this ratio was closer to 1 in V3 than in V1 across both stimulus conditions, indicating that the pRF size difference between achromats and controls was reduced at the higher cortical stage (for the underlying V1 and V3 pRF size by eccentricity distributions, see Supplement 7).

Moreover, if smaller CFs reflect finer sampling of the V1 surface, the V3/V1 pRF-size ratio should itself scale with CF size: the larger the CF size, the more V1 vertices it samples, and therefore the larger the V3/V1 pRF-size ratio should be. Figure 4C shows a vertex-wise analysis across all subjects showing a systematic increase in CF size with V3/V1 pRF-size ratio in both groups and both stimulus conditions. The median rather than the mean was used to summarise the V3/V1 ratio, since some V1 pRF sizes are occasionally close to zero, producing extreme outlier ratios. The relationship between CF and pRF sizes was also present at the subject level: overall, subjects with larger mean CF sizes had larger median V3/V1 ratios (Spearman correlation on log-ratio; non-selective-controls: rho(37)=0.56, p<0.001, achromats: rho(12)=0.60, p=0.023; rod-selective-controls: rho(33)=0.39, p=0.022, achromats: rho(12)=0.40, p=0.159; Figure 4D).

To summarise, we found that achromats’ connective field sizes are smaller than those found in controls, and that connective field size is linked to the size of the V3/V1 pRF-size ratio.

While more work is needed to establish the functional role of this shift and its robustness to confounding factors, we propose it might serve as a compensatory mechanism, where V3’s altered connectivity helps to normalise the processing of altered sensory input from V1.

## Discussion

To test the effects of long-term altered visual input on neural plasticity across the cortical hierarchy, we compared a unique group of individuals who have developed only with rod photoreceptor inputs (achromatopsia) to normally sighted individuals who spend most of their waking time relying on cone-dominated vision. As normally sighted can also experience rod-only vision under specific circumstances, this allowed us to dissociate the effects of life-long altered experience from transient altered sensory input. By integrating markers of plasticity and stability across structural, representational, and connectivity analyses we reveal system-level dynamics of stability and adaptations to altered visual experience. The results offer a new perspective on neural plasticity and stability, recasting these processes as two interacting forces acting simultaneously on the cortex rather than two separate neural responses to change.

Specifically, in achromatopsia we found thickening of the cortex around the foveal projection zone, which receives only cone-photoreceptor inputs and therefore would be signal-deprived from birth in this group. This pattern is consistent with reports of V1 foveal thickening in adult achromats (Molz et al., 2022). Importantly, our sample comprises a substantially younger cohort (8-35 yo) and accounts for age as a covariate, indicating that such structural changes are already present by late childhood. In contrast to early reports (Baseler et al., 2002), we found no evidence that visually-induced signalling inside this deprived region had increased compared to the typically developed rod system. We also observed subtle functional changes in V1 retinotopic tuning including increased population receptive field sizes, however their origins are unclear. These data reveal that visual spatial representations in V1 are largely stable in the face of chronic, highly atypical retinal input, integrating and resolving ambiguity of earlier findings (Baseler et al., 2002; Mckyton et al., 2021; Molz et al., 2022, 2023). This high stability at the cortical input-stage contrasted with reorganisation of the read-out of these inputs by higher-order visual cortex. Notably, more cortical resources in V3 were dedicated to sampling input from the near-fovea, and V3 neuronal populations aggregated information across smaller areas along the V1 cortical surface. There was also evidence for stability of sampling in achromatopsia, because V3 retained connectivity with the foveal confluence despite the life-long scotoma.

Together these findings show that while primary sensory areas such as V1 exhibit considerable stability, higher-order areas such as V3 display marked plasticity that may optimise use of the available information. Both increased sampling of the most fovea-adjacent information available to rods, and finer-grained sampling of coarser inputs could effectively counteract the limitations of developing with a fully rod-based system. However the link between our finding and the potential functional benefit is inferential, and more research is needed to further investigate its origin and function. These findings add a new dimension to the ongoing debate on neural plasticity versus stability by proposing a hierarchical network perspective where both processes interact across processing stages to maintain the integrity of foundational architecture, whilst optimizing the processing of available input.

Several control measures and analyses presented above show that it is unlikely that these results are artefacts of data quality or eye movements. Model fits were comparable across achromat and control groups, ruling out data quality discrepancies as an explanation for the observed differences. Furthermore, converging evidence shows that the lack of ‘filling in’ in the V1 cortex of achromats is unlikely to be obscured by nystagmus (Supplement 2). First, eye-tracking data in achromats showed no correlation between fixation stability and foveal cortex activity. Second, our finding is consistent with two recent studies on adults with achromatopsia who show less severe nystagmus (Kohl et al., 1993), which also found no evidence for filling in (Anderson et al., 2024; Molz et al., 2023). Most crucially, a lack of filling in is aligned with the structural and connectivity analyses revealing corroborating markers of cortical deprivation in line with the pRF mapping result: the deprived V1 foveal projection zone was thickened, and V3’s cortical sampling was shifted away from it.

The hierarchical reorganisation observed in V3 is unlikely to be driven by fixation instability. Connective field (CF) estimates are robust to eye movements (Tangtartharakul et al., 2023), because they are anchored to V1 inputs rather than absolute screen position. Considered alone, the pRF results could alternatively be explained by eye movements introducing a fixed size offset that affects smaller V1 pRFs more strongly than those in V3. While we found no evidence for this relationship between pRF size and gaze measures in our patients, we cannot fully rule out the possibility. Nevertheless, the internal consistency between the CF and pRF measures is in line with the possibility that altered hierarchical sampling contributes to more normalised extrastriate tuning despite coarser tuning at the input stage.

There are convincing arguments for the importance of a system that maintains highly stable sensor-to-cortex connections at the first cortical input stage. Re-routing of cortical projections in response to changing visual signals could lead to “coding catastrophe”, because left unchecked it could leave visual processing areas throughout the brain with mislabelled retinotopic signals, corresponding to different regions of visual space than they would normally represent (Dumoulin & Knapen, 2018; Haak et al., 2015; Wandell & Smirnakis, 2010). Higher-order regions aggregate more inputs across visual space and other sensory domains, and display greater invariance to low-level stimulus properties (Lehky & Tanaka, 2016). Therefore, there is theoretically broader scope for experience-dependent reweighting of inputs (Beyeler et al., 2017; Makin & Krakauer, 2023) and to optimise use of inputs that are still available, more reliable, or more relevant in the impaired system. Conversely, higher-order visual areas may appear more plastic simply because they integrate the cumulative effects of learning from multiple lower stages (Beyeler et al., 2017).

We propose a hierarchical model of neural adaptation that hypothetically could act to improve the limitations of a rod-only visual system, lacking foveal input and acuity. However, to confirm that the adjustments we observe are truly compensatory rather than mere by-products of circuitry responses to visual deprivation, their functional benefits will need to be demonstrated. The adaptations might suggest that individuals with achromatopsia exhibit superior rod-based vision compared to typical systems. One study supporting this, reported improved rod-mediated biological motion processing in some achromatopsia patients, suggesting an enhanced rod system capacity (Burton et al., 2016). Nonetheless, most studies typically find achromatopsia patients’ rod vision to be similar or worse, potentially because rod-related retinal structure can also be impacted by the disease or due to nystagmus (Burton et al., 2016; Hess & Nordby, 1986). It is likely that the functional outcome is shaped by a mixture of impairments and compensatory mechanisms at various levels, which will be important to unpack.

In light of breakthrough treatments set up to restore cone function in human patients with achromatopsia (Farahbakhsh et al., 2022; Fischer et al., 2020; Michaelides et al., 2023; Reichel et al., 2022), the new plasticity dynamics revealed by our work have significant theoretical and practical implications. The possible dormancy of retinotopic maps in cortical input stages as well as some preserved connectivity between V1 foveal areas and V3 bodes well for these gene therapies, each indicating remaining infrastructure to process rescued cone signals. However, our findings also show that the interpretation of signals from V1 by higher-order networks is modified, and it remains to be seen how this affects the translation of new sensory inputs into functional vision. Adaptations in neuronal tuning properties vary in how stably ingrained they become in neural architecture. They can temporarily shift in response to task-demands (transient modulation) (e.g., Klein et al., 2018), or unfold over longer periods with extensive experience that can be acquired or reversed at any age (perceptual learning) (Lu & Dosher, 2022). They can also involve more profound reweighting of neural properties that do not easily revert after critical developmental windows (developmental divergence) (Mitchell & Maurer, 2022). Understanding whether the adaptations in achromatopsia are within the normal range of flexibility for a typically developed system or represent a fundamental reorganisation will be of high relevance to treatment development, as it impacts the nature and extent of neural plasticity that can be harnessed for therapeutic recovery at different ages.

Our findings on hierarchical adaptation have broader implications for other visual disorders, depending on their timing and nature. For instance, a central scotoma acquired in adulthood, as in macular degeneration, may not trigger the same V3 sampling shifts (Haak et al., 2016), suggesting a sensitive window for this form of plasticity, after which connective fields remain more stable. This also raises questions about congenital blindness, where the absence of any driving input could lead to weakening or repurposing of hierarchical connections (Saccone et al., 2024). Moreover, principles may differ between a deprived but structurally intact cortex, as in retinal dystrophies, and a physically damaged cortex, as in stroke. In the latter, more extensive reorganisation may be required to sample effectively from surviving, and potentially disparate, regions of V1. Perceptual training effects in stroke rehabilitation may reflect such dynamics (Cavanaugh et al., 2025; Elshout et al., 2021).

With over 50 interventional studies for inherited retinal diseases initiated, the technology to treat these currently incurable diseases by restoring ocular cellular structure and function is here. The next big hurdle will be to understand neural plasticity constraints on integrating new signals from the eye into the existing neural infrastructure. Our research provides a novel framework for understanding the interactions of neural stability and plasticity as the human brain adapts to altered information. The strength of this approach lies in its use of converging evidence from complementary imaging measures and models. Though each has individual limitations, their combination provides robust support for their underlying mechanisms which are of crucial importance for harnessing the full potential of emerging visual restoration therapies.

## Methods

### Participants

All participants met MRI safety inclusion criteria and had no other known neurological disorders. Patients with achromatopsia had genetically confirmed disease (*CNGA3* or *CNGB3* bi-allelic disease-causing variants). The main dataset was collected at 1.5T. Additional structural data was collected at 3T.

*Main 1.5T dataset:* We obtained functional and structural 1.5T data from fifteen individuals with achromatopsia (age range = 8-27; mean±std = 13.7±5.7), and forty normally sighted controls (age range=6-34; mean±std=19.8±8.3). Additional structural 1.5T data was collected from 8 achromats (age range=8-35; mean±std=19.75±10.6) and 50 normally sighted controls (age range=6-29; mean±std=12.5±5.9).

*Additional 3T dataset (previously published in Anderson et al. 2024):* We obtained additional structural scans from ten normally sighted controls (age range=20-30; mean±std=23.8±3.7) and 9 individuals with achromatopsia (age range=16-33; mean±std=24.2±6.4).

For 4 normally sighted individuals the rod-only pRF map was not collected. One achromat was excluded from functional analysis due to large head-movements. Data collection had ethics approval from the national ethics committee for individuals with achromatopsia (REC reference: 12/LO/1196, IRAS code: 106506, REC reference 11/LO/1229) and the UCL Research Ethics Committee for normally sighted control participants (4846/001).

### MRI acquisition

#### Structural MRI

All 1.5T participants were scanned in a Siemens Avanto 1.5T MRI scanner with a 30-channel coil (a 32-channel coil customised to remove view obstructions). A high-resolution structural scan was acquired using a T1-weighted 3D MPRAGE (1 mm^3^ voxel size, Bandwidth = 190 Hz/Px, 176 partitions, partition TR = 2730, TR = 8.4ms, TE = 3.57, effective T1 = 1000 ms, flip angle = 7 degrees).

All 3T participants were scanned in a Siemens 3T TIM-Trio scanner using a 32-channel head coil. A high-resolution structural scan was acquired using a T1-weighted 3D MDEFT, 1mm isotropic voxels, 176 sagittal slices, 256×240 matrix, TE 2.48ms, TR 7.92ms, TI 910ms.

#### Functional MRI

Functional T2*-weighted multiband 2D echo-planar images were collected using a multi-band sequence with 2.3 mm isotropic voxels (TR = 1000 ms, TE = 55 ms, volumes = 348, flip angle = 75°, MB acceleration factor = 4, Bandwidth = 1628 Hz/Px).

### pRF stimuli and apparatus

Stimuli were presented on an MR-compatible LCD display (BOLDscreen 24, Cambridge Research Systems Ltd., UK; 51 x 32 cm; 1920 x 1200 pixels) viewed through a mirror in the scanner at 105 cm distance. Participants were lying supine in the scanner, with fixation stability recorded where possible via a mirror, with an Eyelink 1000 Plus (SR Research, Ottawa, ON) at the back of bore. Built-in gaze calibration was not possible in achromats due to nystagmus so the camera was calibrated in advance on a healthy eye. Before every other run, patients fixated on a 5-point custom calibration which allowed us to calibrate gaze measures post-hoc (Tailor et al., 2021). Stimuli were presented using MATLAB (R2016b, MathWorks, Natick, MA, USA), via the Psychophysics Toolbox 3 (Kleiner et al., 2007).

pRF mapping stimuli comprised a simultaneous rotating ring and contracting/expanding wedge embedded within a contrast-reversing checkerboard with 2Hz reversal rate. Per run, the ring expanded/contracted for 6 cycles (48 secs/cycle) with logarithmic eccentricity scaling (van Dijk et al., 2016). The wedge (20° angle) rotated clockwise/anticlockwise for 8 cycles (36 sec/cycle). 20-second fixation baselines were embedded at the start, at the mid-point, and end of the run (total run duration 348 secs). The stimuli covered a maximum eccentricity of 8.6°, and moved to a new position each 1-second TR. The stimulus was overlaid with a small white central fixation dot (0.2° VA radius) and a black radial grid to encourage stable fixation. To keep participants engaged, they played a rewarded “kitten rescue mission game” in which they pressed a button when detecting a fixation target luminance change.

The rod-selective (stimulating only rods) and non-selective stimuli (stimulating rods and cones equally) differed in both the colour pair (chromatic pair) used to make up the checkerboard and luminance levels (see Supplement 5). For an in depth description of stimulus generation and validation see Farahbakhsh et al., 2022. In brief, a silent substitution approach was used to obtain a rod-selective colour pair (Estévez & Spekreijse, 1982). We generated a single chromatic pair that kept L and M cone contrast constant (i.e., invisible to L and M cones), whilst inducing a contrast response in rods. Since it is only possible to simultaneously silence two photoreceptor types with 3 colour channels, the resulting stimulus was not controlled for S-cone contribution. To eradicate the S-cone response, we presented these stimuli at very low light level (max luminance: 0.02 cd/m^2^). The non-selective stimuli chromatic pair increased rod and L and M cone contrasts equivalently. This stimulus was presented at mesopic luminance levels (max 0.5 cd/m2).

Prior to the rod-selective condition, which always appeared before the non-selective condition participants lay in the scanner for a 15-minute dark adaptation. A group of 10 normally sighted control adults had a longer period of dark adaptation (45 minutes) to test any impact on the resultant pRF maps. However, since we did not find any differences between the two groups they were combined and treated as a single control group. We note that excluding these 10 participants did not change any of the reported results.

### Fixation stability

Participants’ gaze was tracked throughout all pRF mapping runs. Collecting reliable gaze data from individuals with nystagmus is a challenge because out of the box calibration procedures mostly fail without stable fixation. To account for this, we implemented a post-hoc custom calibration procedure (Tailor et al., 2021). The eye-tracker was first pre-calibrated on a typically sighted individual. Then, before every other run, we collected gaze data from a 5-point fixation task (at fixation and above, below, left, and right of fixation at 5° eccentricity). This data allowed us to subsequently map the patient’s recorded gaze coordinates to their precise locations on the screen. In 10 out of the 14 achromats we acquired reliable enough data to assess fixation stability.

#### Calibration data processing

We first removed the first 0.5 seconds for each fixation location to allow for fixation to arrive on the target. We then performed (a) blink removal, (b) filtered out time points with eye movement velocity outliers (±2SD), and (c) filtered out any positions >3SDs to the left or right of the mean fixation location, and >1SD above or below. We took the median of the remaining gaze measurements as an approximate fixation estimate. The resulting 5 median fixation locations were used to fit an affine transformation that remapped the recorded gaze positions into screen space.

#### Quantifying fixation stability

after applying the transformation of the post-hoc calibration, data was filtered for blinks and extreme velocities (<2SD). For each functional run, fixation instability was measured as the standard deviation of gaze x-positions across 1-second windows. Measures were then averaged across the two run repeats.

### MRI pre-processing

Data were preprocessed using *fMRIPrep* 21.0.2 (Esteban, Blair, et al., 2018; Esteban, Markiewicz, et al., 2018; RRID:SCR_016216), which is based on *Nipype* 1.6.1 (K. Gorgolewski et al., 2011; K. J. Gorgolewski et al., 2018; RRID:SCR_002502).

### fmriprep structural data pre-processing

The T1-weighted (T1w) image was corrected for intensity non-uniformity (INU) with N4BiasFieldCorrection (Tustison et al., 2010), distributed with ANTs 2.3.3 (Avants et al., 2008; RRID:SCR_004757), and used as T1w-reference throughout the workflow. The T1w-reference was then skull-stripped with a *Nipype* implementation of the antsBrainExtraction.sh workflow (from ANTs), using OASIS30ANTs as target template. Brain tissue segmentation of cerebrospinal fluid (CSF), white-matter (WM) and gray-matter (GM) was performed on the brain-extracted T1w using fast (FSL 6.0.5.1:57b01774, RRID:SCR_002823, Zhang et al., 2001). Brain surfaces were reconstructed using recon-all (FreeSurfer 6.0.1, RRID:SCR_001847, Dale et al., 1999), and the brain mask estimated previously was refined with a custom variation of the method to reconcile ANTs-derived and FreeSurfer-derived segmentations of the cortical gray-matter of Mindboggle (RRID:SCR_002438, A. Klein et al., 2017). Volume-based spatial normalization to one standard space (MNI152NLin2009cAsym) was performed through nonlinear registration with antsRegistration (ANTs 2.3.3), using brain-extracted versions of both T1w reference and the T1w template. The following template was selected for spatial normalization: *ICBM 152 Nonlinear Asymmetrical template version 2009c* [Fonov et al., 2009, RRID:SCR_008796; TemplateFlow ID: MNI152NLin2009cAsym]. Many internal operations of ***fMRIPrep*** use ***Nilearn*** 0.8.1 (Abraham et al., 2014, RRID:SCR_001362), mostly within the functional processing workflow.

### fmriprep functional data pre-processing

For each BOLD run found per subject (across all tasks), the following preprocessing was performed. First, a reference volume and its skull-stripped version were generated by aligning and averaging 1 single-band references (SBRefs). Head-motion parameters with respect to the BOLD reference (transformation matrices, and six corresponding rotation and translation parameters) are estimated before any spatiotemporal filtering using mcflirt (FSL 6.0.5.1:57b01774, Jenkinson et al., 2002). The BOLD time-series were resampled onto their original, native space by applying the transforms to correct for head-motion. These resampled BOLD time-series will be referred to as *preprocessed BOLD in original space*, or just *preprocessed BOLD*. The BOLD reference was then co-registered to the T1w reference using bbregister (FreeSurfer) which implements boundary-based registration (Greve & Fischl, 2009). Co-registration was configured with six degrees of freedom. First, a reference volume and its skull-stripped version were generated using a custom methodology of *fMRIPrep*. Several confounding time-series were calculated based on the *preprocessed BOLD*: framewise displacement (FD), DVARS and three region-wise global signals. FD was computed using two formulations following Power (absolute sum of relative motions, Power et al., 2014) and Jenkinson (relative root mean square displacement between affines, Jenkinson et al., 2002). FD and DVARS are calculated for each functional run, both using their implementations in *Nipype* (following the definitions by Power et al., 2014). The three global signals are extracted within the CSF, the WM, and the whole-brain masks. Additionally, a set of physiological regressors were extracted to allow for component-based noise correction (*CompCor*, Behzadi et al., 2007). Principal components are estimated after high-pass filtering the *preprocessed BOLD* time-series (using a discrete cosine filter with 128s cut-off) for the two *CompCor* variants: temporal (tCompCor) and anatomical (aCompCor). tCompCor components are then calculated from the top 2% variable voxels within the brain mask. For aCompCor, three probabilistic masks (CSF, WM and combined CSF+WM) are generated in anatomical space. The implementation differs from that of Behzadi et al. in that instead of eroding the masks by 2 pixels on BOLD space, the aCompCor masks are subtracted a mask of pixels that likely contain a volume fraction of GM. This mask is obtained by dilating a GM mask extracted from the FreeSurfer’s *aseg* segmentation, and it ensures components are not extracted from voxels containing a minimal fraction of GM. Finally, these masks are resampled into BOLD space and binarized by thresholding at 0.99 (as in the original implementation). Components are also calculated separately within the WM and CSF masks. For each CompCor decomposition, the *k* components with the largest singular values are retained, such that the retained components’ time series are sufficient to explain 50 percent of variance across the nuisance mask (CSF, WM, combined, or temporal). The remaining components are dropped from consideration. The head-motion estimates calculated in the correction step were also placed within the corresponding confounds file. The confound time series derived from head motion estimates and global signals were expanded with the inclusion of temporal derivatives and quadratic terms for each (Satterthwaite et al., 2013). Frames that exceeded a threshold of 0.9 mm FD or 1.5 standardised DVARS were annotated as motion outliers. The BOLD time-series were resampled onto the following surfaces (FreeSurfer reconstruction nomenclature): *fsnative*. All resamplings can be performed with *a single interpolation step* by composing all the pertinent transformations (i.e., head-motion transform matrices, susceptibility distortion correction when available, and co-registrations to anatomical and output spaces). Surface resampling was performed using mri_vol2surf (FreeSurfer).

### Additional pre-processing

The first 4 volumes of each run were discarded for signal stabilization. Bold data in fsnative space was smoothed along the cortical surface (Gaussian kernel FWHM = 3 mm). For each run, a denoising procedure was further applied on the data by inputting nuisance regressors into a PCA and regressing out the top 5 components. The nuisance regressors include: Six head motion regressors, FD and DVARS, CSF mean signal, white matter mean signal, the first five anatomical CompCor regressors (csf+wm combined), and the first five temporal CompCor regressors. Then data and regressors were temporally filtered with a Savitzky-Golay filter (3rd order, 347 seconds in length) and zscored. The time series from the two identical task runs were then averaged to increase signal to noise ratio.

### MRI analysis

#### Region of interests

An anatomically defined retinotopy atlas was applied to the inflated cortical surface for each participant in order to delineate the borders of V1, V2 and V3, and also each degree of eccentricity up to 8° in V1 (Benson & Winawer, 2018). See Figure 1B for an example of an eccentricity plot for one participant.

#### Structural data analysis

The ‘mris_anatomical_stats’ command was used to extract surface area and cortical thickness properties for each participant, per hemisphere, for each region of interest. Overall cortical thickness per participant was computed by weighting the cortical thickness of a hemisphere by the number of vertices per hemisphere.

#### pRF analysis

pRF modelling was carried out with a custom Python package, prfpy (Aqil & Knapen, 2023). To model population receptive fields, we used an isometric bivariate Gaussian model (Dumoulin & Wandell, 2008), with *x*,*y* representing the preferred retinotopic position, spatial standard deviation sigma (σ) representing pRF size as well as the baseline of the timeseries and the signal amplitude beta (β). At every cortical location, the parameters of each model were optimized by minimizing the residual sum of squares between the model prediction and the measured BOLD signal. Fitting began with a grid search for the best model parameters (pRF position and size) followed by an iterative optimization. In each participant, in each condition, vertices with an R^2^ < 0.1 (variance explained by the pRF model), a negative beta, or a centre outside the stimulated radius (8.6°) were excluded from further analysis.

In addition, we ran a second model fit using a stimulus aperture file that incorporated a black oval mask (1.25°x0.96°) over the area of the central rod-free scotoma (Curcio et al., 1990). Previous research suggests that biases in pRF measures such as shifts in eccentricities and size can occur when a scotoma is present, and can be partially accounted for when it is explicitly modelled in the stimulus apertures (Binda et al., 2013). Results from this analysis are reported in Supplement 3.

#### Connective field modelling

Connective field modelling was carried out with a dedicated Python package, prfpy (Aqil & Knapen, 2023) based on the method described in Knapen (2021). In brief, gaussian connective field profiles on the surface are defined for each vertex on the cortical manifold, with a centre location of the connective field (v-centre), and σ (sigma) the Gaussian spread in millimeters on the cortical surface.

To generate spatial Gaussian fields along the V1 surface, the diffusion of heat along the cortical mesh was simulated (Crane et al., 2013) and used to infer geodesic distances between all V1 vertices (V1 definition was based on the Benson atlas, see Regions of Interest). As the two hemispheres are two separate surface meshes, this distance matrix was calculated for each hemisphere separately. Grid-fit candidate CF sizes were: [0.5, 1, 2, 3, 4, 5, 7, 10, 15, 20] mm σ for the Gaussian CF. The predicted time course for each of the resulting CF models was generated by taking the dot product between the CF’s vertex profile and the vertex by time-courses matrix in V1. The resulting CF model time courses were z scored and correlated with the time courses throughout the brain. For each location in the brain, the connective field resulting in the highest squared correlation was selected. Only values within an R^2^ > 0.1, beta > 0, and a centre within 8° of eccentricity (matching the radius of the stimuli) were included in further analyses. To create connective-field-based eccentricity maps, each vertex was assigned the Benson atlas-based eccentricities (Benson & Winawer, 2018) of its CF V1 centre.

To relate CF size to spatial tuning in V3, we computed a V3/V1 pRF-size ratio for each V3 vertex. Each V3 vertex’s pRF size was divided by the pRF size of the V1 vertex at its CF centre (v-centre), estimated from the same stimulus condition. Only V3 vertices that met both the pRF and CF inclusion criteria, and whose V1 centre vertex met the pRF inclusion criteria, were included. Ratios were summarised using the median rather than the mean, as V1 pRF sizes close to zero occasionally produced extreme ratios. For the vertex-wise analysis, V3 vertices were pooled across participants within each group and condition. The median ratio and interquartile range were then computed for each CF size (Figure 4C). For the subject-level analysis, we computed each participant’s median V3/V1 ratio and mean CF size. The ratio was log-transformed, and its relationship with CF size was assessed using Spearman correlations (Figure 4D).

For visualisation of group maps all individual maps were transformed to fsaverage space. Plotted values indicate the medians of each group and condition per vertex. A vertex was left empty if no significant data was available for more than 30% of individuals within the group.

### Statistical analysis

Jamovi R package (jmv) was used for Anovas and t-tests (The jamovi project, 2022). Jasp (jsq) R package for Bayesian t-tests (JASP team, 2020). When sphericity assumptions were not met a greenhouse-geisser correction was applied. Uncorrected p-values are reported as well as FDR multiple comparison corrections. Spearman rank correlations were used to assess subject-level relationships between CF size and the V3/V1 pRF-size ratio. V3/V1 pRF size ratios were log-transformed as the ratio is a multiplicative quantity referenced to 1.

## Acknowledgments

We thank Maria Molina Sanchez, Hunter Schone and Matan Mazor for providing comments on an earlier version of this manuscript.

## References

Aboshiha, J., Dubis, A. M., Cowing, J., Fahy, R. T. A., Sundaram, V., Bainbridge, J. W., Ali, R. R., Dubra, A., Nardini, M., Webster, A. R., Moore, A. T., Rubin, G., Carroll, J., & Michaelides, M. (2014). A prospective longitudinal study of retinal structure and function in achromatopsia. Investigative Ophthalmology and Visual Science, 55(9), 5733–5743. 10.1167/iovs.14-14937

Abraham, A., Pedregosa, F., Eickenberg, M., Gervais, P., Mueller, A., Kossaifi, J., Gramfort, A., Thirion, B., & Varoquaux, G. (2014). Machine learning for neuroimaging with scikit-learn. Frontiers in Neuroinformatics, 8. 10.3389/fninf.2014.00014

Adams, D. L., Sincich, L. C., & Horton, J. C. (2007). Complete Pattern of Ocular Dominance Columns in Human Primary Visual Cortex. Journal of Neuroscience, 27(39), 10391–10403. 10.1523/JNEUROSCI.2923-07.2007

Anderson, E. J., Dekker, T. M., Farahbakhsh, M., Hirji, N., Schwarzkopf, D. S., Michaelides, M., & Rees, G. (2024). fMRI and gene therapy in adults with CNGB3 mutation. Brain Research Bulletin, 215, 111026. 10.1016/j.brainresbull.2024.111026

Aqil, M., & Knapen, T. (2023). Prfpy: A python package to simulate and fit population receptive field models to time series data. (Version v0.1.0-alpha) [Computer software]. Zenodo. 10.5281/zenodo.10201022

Avants, B. B., Epstein, C. L., Grossman, M., & Gee, J. C. (2008). Symmetric diffeomorphic image registration with cross-correlation: Evaluating automated labeling of elderly and neurodegenerative brain. Medical Image Analysis, 12(1), 26–41. 10.1016/j.media.2007.06.004

Baker, C. I., Dilks, D. D., Peli, E., & Kanwisher, N. (2008). Reorganization of visual processing in macular degeneration: Replication and clues about the role of foveal loss. Vision Research, 48(18), 1910–1919. 10.1016/j.visres.2008.05.020

Baker, C. I., Peli, E., Knouf, N., & Kanwisher, N. G. (2005). Reorganization of Visual Processing in Macular Degeneration. The Journal of Neuroscience, 25(3), 614–618. 10.1523/JNEUROSCI.3476-04.2005

Barton, B., & Brewer, A. A. (2015). fMRI of the rod scotoma elucidates cortical rod pathways and implications for lesion measurements. Proceedings of the National Academy of Sciences of the United States of America, 112(16), 5201–5206. 10.1073/PNAS.1423673112/-/DCSUPPLEMENTAL

Baseler, H. A., Brewer, A. A., Sharpe, L. T., Morland, A. B., Jaägle, H., & Wandell, B. A. (2002). Reorganization of human cortical maps caused by inherited photoreceptor abnormalities. Nature Neuroscience 2002 5:4, 5(4), 364–370. 10.1038/nn817

Baseler, H. A., Gouws, A., Haak, K. V., Racey, C., Crossland, M. D., Tufail, A., Rubin, G. S., Cornelissen, F. W., & Morland, A. B. (2011). Large-scale remapping of visual cortex is absent in adult humans with macular degeneration. Nature Neuroscience, 14(5), 649–657. 10.1038/nn.2793

Behzadi, Y., Restom, K., Liau, J., & Liu, T. T. (2007). A component based noise correction method (CompCor) for BOLD and perfusion based fMRI. NeuroImage, 37(1), 90–101. 10.1016/j.neuroimage.2007.04.042

Benson, N. C., & Winawer, J. (2018). Bayesian analysis of retinotopic maps. eLife, 7. 10.7554/ELIFE.40224

Beyeler, M., Rokem, A., Boynton, G. M., & Fine, I. (2017). Learning to see again: Biological constraints on cortical plasticity and the implications for sight restoration technologies. Journal of Neural Engineering, 14(5), 051003. 10.1088/1741-2552/aa795e

Binda, P., Thomas, J. M., Boynton, G. M., & Fine, I. (2013). Minimizing biases in estimating the reorganization of human visual areas with BOLD retinotopic mapping. Journal of Vision, 13(7), 13. 10.1167/13.7.13

Brown, H. D. H., Gouws, A. D., Vernon, R. J. W., Lawrence, S. J. D., Donnelly, G., Gill, L., Gale, R. P., Baseler, H. A., & Morland, A. B. (2021). Assessing functional reorganization in visual cortex with simulated retinal lesions. Brain Structure and Function, 226(9), 2855–2867. 10.1007/s00429-021-02366-w

Burton, E., Wattam-Bell, J., S. Rubin, G., Aboshiha, J., Michaelides, M., Atkinson, J., Braddick, O., & Nardini, M. (2016). Dissociations in Coherence Sensitivity Reveal Atypical Development of Cortical Visual Processing in Congenital Achromatopsia. Investigative Ophthalmology & Visual Science, 57(4), 2251–2259. 10.1167/iovs.15-18414

Calford, M. B., Chino, Y. M., Das, A., Eysel, U. T., Gilbert, C. D., Heinen, S. J., Kaas, J. H., & Ullman, S. (2005). Rewiring the adult brain. Nature, 438(7065), E3–E3. 10.1038/nature04359

Carvalho, J., Renken, R. J., & Cornelissen, F. W. (2019). Studying Cortical Plasticity in Ophthalmic and Neurological Disorders: From Stimulus-Driven to Cortical Circuitry Modeling Approaches. Neural Plasticity, 2019, e2724101. 10.1155/2019/2724101

Carvalho, J., Renken, R. J., & Cornelissen, F. W. (2021). Predictive masking of an artificial scotoma is associated with a system-wide reconfiguration of neural populations in the human visual cortex. NeuroImage, 245, 118690. 10.1016/j.neuroimage.2021.118690

Cavanaugh, M. R., Fahrenthold, B. K., & Huxlin, K. R. (2025). What V1 Damage Can Teach Us About Visual Perception and Learning. Annual Review of Vision Science, 11(1), 217–241. 10.1146/annurev-vision-110323-112823

Crane, K., Weischedel, C., & Wardetzky, M. (2013). Geodesics in heat: A new approach to computing distance based on heat flow. ACM Transactions on Graphics, 32(5), 152:1–152:11. 10.1145/2516971.2516977

Curcio, C. A., Sloan, K. R., Kalina, R. E., & Hendrickson, A. E. (1990). Human photoreceptor topography. Journal of Comparative Neurology, 292(4), 497–523. 10.1002/cne.902920402

Dale, A. M., Fischl, B., & Sereno, M. I. (1999). Cortical Surface-Based Analysis: I. Segmentation and Surface Reconstruction. NeuroImage, 9(2), 179–194. 10.1006/nimg.1998.0395

Dilks, D. D., Baker, C. I., Peli, E., & Kanwisher, N. (2009). Reorganization of Visual Processing in Macular Degeneration Is Not Specific to the “Preferred Retinal Locus”. The Journal of Neuroscience, 29(9), 2768. 10.1523/JNEUROSCI.5258-08.2009

Dumoulin, S. O., & Knapen, T. (2018). How Visual Cortical Organization Is Altered by Ophthalmologic and Neurologic Disorders. Annual Review of Vision Science, 4(1), 357–379. 10.1146/annurev-vision-091517-033948

Dumoulin, S. O., & Wandell, B. A. (2008). Population receptive field estimates in human visual cortex. NeuroImage, 39(2), 647–660. 10.1016/j.neuroimage.2007.09.034

Elshout, J. A., Bergsma, D. P., van den Berg, A. V., & Haak, K. V. (2021). Functional MRI of visual cortex predicts training-induced recovery in stroke patients with homonymous visual field defects. NeuroImage: Clinical, 31, 102703. 10.1016/j.nicl.2021.102703

Esteban, O., Blair, R., Markiewicz, C. J., Berleant, S. L., Moodie, C., Ma, F., Isik, A. I., Erramuzpe, A., Kent, M., James D. and Goncalves, DuPre, E., Sitek, K. R., Gomez, D. E. P., Lurie, D. J., Ye, Z., Poldrack, R. A., & Gorgolewski, K. J. (2018). fMRIPrep. Software. 10.5281/zenodo.852659

Esteban, O., Markiewicz, C., Blair, R. W., Moodie, C., Isik, A. I., Erramuzpe Aliaga, A., Kent, J., Goncalves, M., DuPre, E., Snyder, M., Oya, H., Ghosh, S., Wright, J., Durnez, J., Poldrack, R., & Gorgolewski, K. J. (2018). fMRIPrep: A robust preprocessing pipeline for functional MRI. Nature Methods. 10.1038/s41592-018-0235-4

Estévez, O., & Spekreijse, H. (1982). The ‘silent substitution’ method in visual research. Vision Research, 22(6), 681–691. 10.1016/0042-6989(82)90104-3

Farahbakhsh, M., Anderson, E. J., Maimon-Mor, R. O., Rider, A., Greenwood, J. A., Hirji, N., Zaman, S., Jones, P. R., Schwarzkopf, D. S., Rees, G., Michaelides, M., & Dekker, T. M. (2022). A demonstration of cone function plasticity after gene therapy in achromatopsia. Brain, 145(11), 3803–3815. 10.1093/brain/awac226

Ferreira, S., Pereira, A. C., Quendera, B., Reis, A., Silva, E. D., & Castelo-Branco, M. (2016). Primary visual cortical remapping in patients with inherited peripheral retinal degeneration. NeuroImage : Clinical, 13, 428–438. 10.1016/j.nicl.2016.12.013

Fischer, M. D., Michalakis, S., Wilhelm, B., Zobor, D., Muehlfriedel, R., Kohl, S., Weisschuh, N., Ochakovski, G. A., Klein, R., Schoen, C., Sothilingam, V., Garcia-Garrido, M., Kuehlewein, L., Kahle, N., Werner, A., Dauletbekov, D., Paquet-Durand, F., Tsang, S., Martus, P., … Wissinger, B. (2020). Safety and Vision Outcomes of Subretinal Gene Therapy Targeting Cone Photoreceptors in Achromatopsia: A Nonrandomized Controlled Trial. JAMA Ophthalmology, 138(6), 643–651. 10.1001/jamaophthalmol.2020.1032

Fonov, V., Evans, A., McKinstry, R., Almli, C., & Collins, D. (2009). Unbiased nonlinear average age-appropriate brain templates from birth to adulthood. NeuroImage, 47, Supplement 1, S102. 10.1016/S1053-8119(09)70884-5

Georgiou, M., Fujinami, K., & Michaelides, M. (2021). Inherited retinal diseases: Therapeutics, clinical trials and end points—A review. Clinical & Experimental Ophthalmology, 49(3), 270–288. 10.1111/ceo.13917

Goesaert, E., Van Baelen, M., Spileers, W., Wagemans, J., & Op de Beeck, H. P. (2014). Visual Space and Object Space in the Cerebral Cortex of Retinal Disease Patients. PLoS ONE, 9(2), e88248. 10.1371/journal.pone.0088248

Gorgolewski, K., Burns, C. D., Madison, C., Clark, D., Halchenko, Y. O., Waskom, M. L., & Ghosh, S. (2011). Nipype: A flexible, lightweight and extensible neuroimaging data processing framework in Python. Frontiers in Neuroinformatics, 5, 13. 10.3389/fninf.2011.00013

Gorgolewski, K. J., Esteban, O., Markiewicz, C. J., Ziegler, E., Ellis, D. G., Notter, M. P., Jarecka, D., Johnson, H., Burns, C., Manhães-Savio, A., Hamalainen, C., Yvernault, B., Salo, T., Jordan, K., Goncalves, M., Waskom, M., Clark, D., Wong, J., Loney, F., … Ghosh, S. (2018). Nipype. Software. 10.5281/zenodo.596855

Greve, D. N., & Fischl, B. (2009). Accurate and robust brain image alignment using boundary-based registration. NeuroImage, 48(1), 63–72. 10.1016/j.neuroimage.2009.06.060

Haak, K. V., Cornelissen, F. W., & Morland, A. B. (2012). Population Receptive Field Dynamics in Human Visual Cortex. PLoS ONE, 7(5), e37686. 10.1371/journal.pone.0037686

Haak, K. V., Morland, A. B., & Engel, S. A. (2015). Plasticity, and its limits, in adult human primary visual cortex. Multisensory Research, 28(3–4), 297–307. 10.1163/22134808-00002496

Haak, K. V., Morland, A. B., Rubin, G. S., & Cornelissen, F. W. (2016). Preserved retinotopic brain connectivity in macular degeneration. Ophthalmic and Physiological Optics, 36(3), 335–343. 10.1111/opo.12279

Haak, K. V., Winawer, J., Harvey, B. M., Renken, R., Dumoulin, S. O., Wandell, B. A., & Cornelissen, F. W. (2013). Connective field modeling. NeuroImage, 66, 376–384. 10.1016/j.neuroimage.2012.10.037

Hess, R. F., & Nordby, K. (1986). Spatial and temporal limits of vision in the achromat. The Journal of Physiology, 371(1), 365–385. 10.1113/jphysiol.1986.sp015981

Hummer, A., Ritter, M., Woletz, M., Ledolter, A. A., Tik, M., Dumoulin, S. O., Holder, G. E., Schmidt-Erfurth, U., & Windischberger, C. (2018). Artificial scotoma estimation based on population receptive field mapping. NeuroImage, 169, 342–351. 10.1016/J.NEUROIMAGE.2017.12.010

JASP team. (2020). JASP (Version 0.18.3) [Computer software]. https://jasp-stats.org/

Jenkinson, M., Bannister, P., Brady, M., & Smith, S. (2002). Improved Optimization for the Robust and Accurate Linear Registration and Motion Correction of Brain Images. NeuroImage, 17(2), 825–841. 10.1006/nimg.2002.1132

Jiang, J., Zhu, W., Shi, F., Liu, Y., Li, J., Qin, W., Li, K., Yu, C., & Jiang, T. (2009). Thick Visual Cortex in the Early Blind. Journal of Neuroscience, 29(7), 2205–2211. 10.1523/JNEUROSCI.5451-08.2009

Jones, P. R., Somoskeöy, T., Chow-Wing-Bom, H., & Crabb, D. P. (2020). Seeing other perspectives: Evaluating the use of virtual and augmented reality to simulate visual impairments (OpenVisSim). Npj Digital Medicine, 3(1), 1–9. 10.1038/s41746-020-0242-6

Klein, A., Ghosh, S. S., Bao, F. S., Giard, J., Häme, Y., Stavsky, E., Lee, N., Rossa, B., Reuter, M., Neto, E. C., & Keshavan, A. (2017). Mindboggling morphometry of human brains. PLOS Computational Biology, 13(2), e1005350. 10.1371/journal.pcbi.1005350

Klein, B. P., Fracasso, A., van Dijk, J. A., Paffen, C. L. E., te Pas, S. F., & Dumoulin, S. O. (2018). Cortical depth dependent population receptive field attraction by spatial attention in human V1. NeuroImage, 176, 301–312. 10.1016/j.neuroimage.2018.04.055

Kleiner, M., Brainard, D. H., Pelli, D. G., Broussard, C., Wolf, T., & Niehorster, D. (2007). What’s new in Psychtoolbox-3? A free cross-platform toolkit for psychophysiscs with Matlab and GNU/Octave. In Cognitive and Computational Psychophysics (Vol. 36).

Knapen, T. (2021). Topographic connectivity reveals task-dependent retinotopic processing throughout the human brain. Proceedings of the National Academy of Sciences, 118(2), e2017032118. 10.1073/pnas.2017032118

Kohl, S., Jägle, H., Wissinger, B., & Zobor, D. (1993). Achromatopsia. In M. P. Adam, J. Feldman, G. M. Mirzaa, R. A. Pagon, S. E. Wallace, & A. Amemiya (Eds), GeneReviews®. University of Washington, Seattle. http://www.ncbi.nlm.nih.gov/books/NBK1418/

Lehky, S. R., & Tanaka, K. (2016). Neural representation for object recognition in inferotemporal cortex. Current Opinion in Neurobiology, Neurobiology of Cognitive Behavior, 37, 23–35. 10.1016/j.conb.2015.12.001

Lu, Z.-L., & Dosher, B. A. (2022). Current directions in visual perceptual learning. Nature Reviews Psychology, 1(11), 654–668. 10.1038/s44159-022-00107-2

Macnamara, A., Chen, C., Schinazi, V. R., Saredakis, D., & Loetscher, T. (2021). Simulating Macular Degeneration to Investigate Activities of Daily Living: A Systematic Review. Frontiers in Neuroscience, 15. 10.3389/fnins.2021.663062

Makin, T. R., & Krakauer, J. W. (2023). Against cortical reorganisation. eLife, 12, e84716. 10.7554/eLife.84716

Masuda, Y., Takemura, H., Terao, M., Miyazaki, A., Ogawa, S., Horiguchi, H., Nakadomari, S., Matsumoto, K., Nakano, T., Wandell, B. A., & Amano, K. (2021). V1 Projection Zone Signals in Human Macular Degeneration Depend on Task Despite Absence of Visual Stimulus. Current Biology, 31(2), 406–412.e3. 10.1016/j.cub.2020.10.034

Mckyton, A., Averbukh, E., Marks Ohana, D., Levin, N., & Banin, E. (2021). Cortical Visual Mapping following Ocular Gene Augmentation Therapy for Achromatopsia. 10.1523/JNEUROSCI.3222-20.2021

Michaelides, M., Hirji, N., Wong, S. C., Besirli, C. G., Zaman, S., Kumaran, N., Georgiadis, A., Smith, A. J., Ripamonti, C., Gottlob, I., Robson, A. G., Thiadens, A., Henderson, R. H., Fleck, P., Anglade, E., Dong, X., Capuano, G., Lu, W., Berry, P., … Bainbridge, J. (2023). First-in-Human Gene Therapy Trial of AAV8-hCARp.hCNGB3 in Adults and Children With CNGB3-associated Achromatopsia. American Journal of Ophthalmology, 253, 243–251. 10.1016/j.ajo.2023.05.009

Mitchell, D. E., & Maurer, D. (2022). Critical Periods in Vision Revisited. Annual Review of Vision Science, 8(1), 291–321. 10.1146/annurev-vision-090721-110411

Molz, B., Herbik, A., Baseler, H. A., de Best, P. B., Vernon, R. W., Raz, N., Gouws, A. D., Ahmadi, K., Lowndes, R., McLean, R. J., Gottlob, I., Kohl, S., Choritz, L., Maguire, J., Kanowski, M., Käsmann-Kellner, B., Wieland, I., Banin, E., Levin, N., … Morland, A. B. (2022). Structural changes to primary visual cortex in the congenital absence of cone input in achromatopsia. NeuroImage: Clinical, 33, 102925. 10.1016/J.NICL.2021.102925

Molz, B., Herbik, A., Baseler, H. A., de Best, P., Raz, N., Gouws, A., Ahmadi, K., Lowndes, R., McLean, R. J., Gottlob, I., Kohl, S., Choritz, L., Maguire, J., Kanowski, M., Käsmann-Kellner, B., Wieland, I., Banin, E., Levin, N., Morland, A. B., & Hoffmann, M. B. (2023). Achromatopsia—Visual Cortex Stability and Plasticity in the Absence of Functional Cones. Investigative Ophthalmology & Visual Science, 64(13), 23. 10.1167/iovs.64.13.23

Park, H.-J., Lee, J. D., Kim, E. Y., Park, B., Oh, M.-K., Lee, S., & Kim, J.-J. (2009). Morphological alterations in the congenital blind based on the analysis of cortical thickness and surface area. NeuroImage, 47(1), 98–106. 10.1016/j.neuroimage.2009.03.076

Power, J. D., Mitra, A., Laumann, T. O., Snyder, A. Z., Schlaggar, B. L., & Petersen, S. E. (2014). Methods to detect, characterize, and remove motion artifact in resting state fMRI. NeuroImage, 84(Supplement C), 320–341. 10.1016/j.neuroimage.2013.08.048

Reichel, F. F., Michalakis, S., Wilhelm, B., Zobor, D., Muehlfriedel, R., Kohl, S., Weisschuh, N., Sothilingam, V., Kuehlewein, L., Kahle, N., Seitz, I., Paquet-Durand, F., Tsang, S. H., Martus, P., Peters, T., Seeliger, M., Bartz-Schmidt, K. U., Ueffing, M., Zrenner, E., … Fischer, D. (2022). Three-year results of phase I retinal gene therapy trial for CNGA3-mutated achromatopsia: Results of a non randomised controlled trial. British Journal of Ophthalmology, 106(11), 1567–1572. 10.1136/bjophthalmol-2021-319067

Ritter, M., Hummer, A., Ledolter, A. A., Holder, G. E., Windischberger, C., & Schmidt-Erfurth, U. M. (2019). Correspondence between retinotopic cortical mapping and conventional functional and morphological assessment of retinal disease. British Journal of Ophthalmology, 103(2), 208–215. 10.1136/bjophthalmol-2017-311443

Saccone, E. J., Tian, M., & Bedny, M. (2024). Developing cortex is functionally pluripotent: Evidence from blindness. Developmental Cognitive Neuroscience, 66, 101360. 10.1016/j.dcn.2024.101360

Satterthwaite, T. D., Elliott, M. A., Gerraty, R. T., Ruparel, K., Loughead, J., Calkins, M. E., Eickhoff, S. B., Hakonarson, H., Gur, R. C., Gur, R. E., & Wolf, D. H. (2013). An improved framework for confound regression and filtering for control of motion artifact in the preprocessing of resting-state functional connectivity data. NeuroImage, 64(1), 240–256. 10.1016/j.neuroimage.2012.08.052

Schumacher, E. H., Jacko, J. A., Primo, S. A., Main, K. L., Moloney, K. P., Kinzel, E. N., & Ginn, J. (2008). Reorganization of visual processing is related to eccentric viewing in patients with macular degeneration. Restorative Neurology and Neuroscience, 26(4– 5), 391–402.

Tailor, V. K., Theodorou, M., Dahlmann-Noor, A. H., Dekker, T. M., & Greenwood, J. A. (2021). Eye movements elevate crowding in idiopathic infantile nystagmus syndrome. Journal of Vision, 21(13), 9–9. 10.1167/JOV.21.13.9

Tangtartharakul, G., Morgan, C. A., Rushton, S. K., & Schwarzkopf, D. S. (2023). Retinotopic connectivity maps of human visual cortex with unconstrained eye movements. Human Brain Mapping, 44(16), 5221–5237. 10.1002/hbm.26446

The jamovi project. (2022). *Jamovi* (Version 2.3) [Computer software]. https://www.jamovi.org

Tustison, N. J., Avants, B. B., Cook, P. A., Zheng, Y., Egan, A., Yushkevich, P. A., & Gee, J. C. (2010). N4ITK: Improved N3 Bias Correction. IEEE Transactions on Medical Imaging, 29(6), 1310–1320. 10.1109/TMI.2010.2046908

van Dijk, J. A., de Haas, B., Moutsiana, C., & Schwarzkopf, D. S. (2016). Intersession reliability of population receptive field estimates. NeuroImage, 143, 293–303. 10.1016/j.neuroimage.2016.09.013

Wandell, B. A., & Smirnakis, S. M. (2010). Plasticity and stability of visual field maps in adult primary visual cortex. Nature Reviews Neuroscience, 10, 873–884. 10.1038/nrn2741

Zhang, Y., Brady, M., & Smith, S. (2001). Segmentation of brain MR images through a hidden Markov random field model and the expectation-maximization algorithm. IEEE Transactions on Medical Imaging, 20(1), 45–57. 10.1109/42.906424

