## Supplementary Materials for "Hierarchical cortical plasticity in congenital sight impairment"

#### Supplementary Material 1. High foveal coverage achromat individuals

##### Eccentricity

##### Polar angle

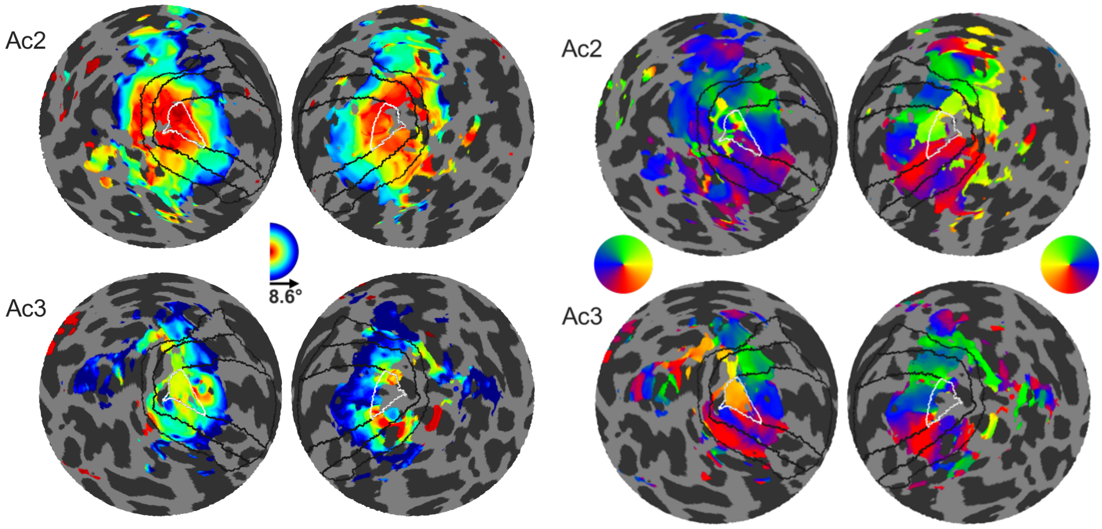


*Figure S1. Eccentricity and polar maps of both hemispheres in the two achromats that fall outside the rod-selective control 95% predictive interval. Maps are derived from activity in the non-selective stimulation condition. Black lines denote delineations of V1,2,3; white lines mark the foveal confluence in V1.*

AC2 (top) has a control-like map with foveal eccentricities being represented in the foveal projection zone (0°-2°) in V1. This means foveal input is coming in from the retina into the foveal projection zone and would not be considered a form of remapping. This could be explained by either a presence of rods in the usually rod-free retinal zone or an indication of residual cone-function. AC3 (bottom) shows peripheral activations in the foveal projection zone of their left hemisphere. However, the polar angle map in the foveal projection zone does not follow the usual retinotopic pattern of upper to lower visual field, in this visualisation red to green.

### Supplementary Material 2. Fixation stability

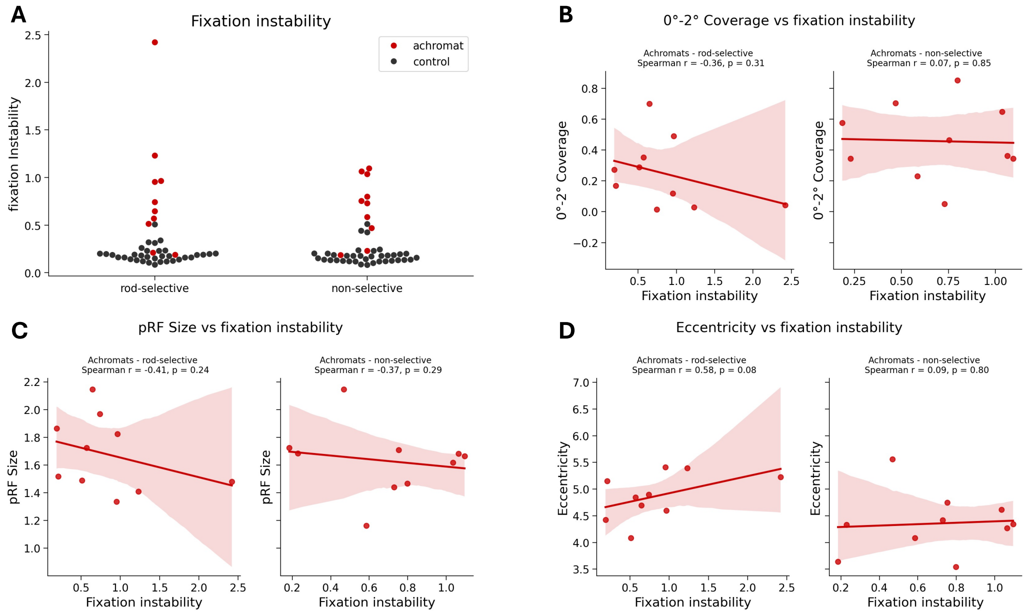


*Figure S2A. (A) Fixation instability measures across both groups in each stimulation condition. (B) Relationship between coverage in the foveal projection zone and fixation instability (C) Relationship between pRF size (across V1) and fixation instability (D) Relationship between eccentricity (across V1) and fixation instability.*

As expected, achromats showed significantly higher fixation instability compared to controls (as reported in the main text). We found no significant correlation between fixation instability and either coverage, pRF size, eccentricity in achromats. Results of Spearman R correlations in both rod- and non-selective conditions are reported in the figure. We note that the relationship between nystagmus and visual sampling is complex - patients experience a stable image and may sample only during specific eye-movement phases. It is therefore not fully clear if and how nystagmus should give rise to altered pRFs.

Gaze data could not be analysed for four achromats. For one participant, no data were recorded due to a technical issue with the eye tracker. For two, head position in the scanner with respect to the eye tracker left too few usable samples. For one, high gaze variability during post-hoc calibration meant the five fixation points could not be distinguished. To test whether excluding these participants biased the results, we repeated the correlations with all four assigned a fixation instability value of 3. This is higher than the least stable calibrated participant. The results were unchanged. Fixation instability did not correlate with pRF size (rod-selective: Spearman $r(12) = -0.31$, $p = 0.28$; non-selective: $r(12) = -0.34$, $p = 0.24$) or eccentricity (rod-selective: $r(12) = -0.08$, $p = 0.80$; non-selective: $r(12) = -0.10$, $p = 0.73$).

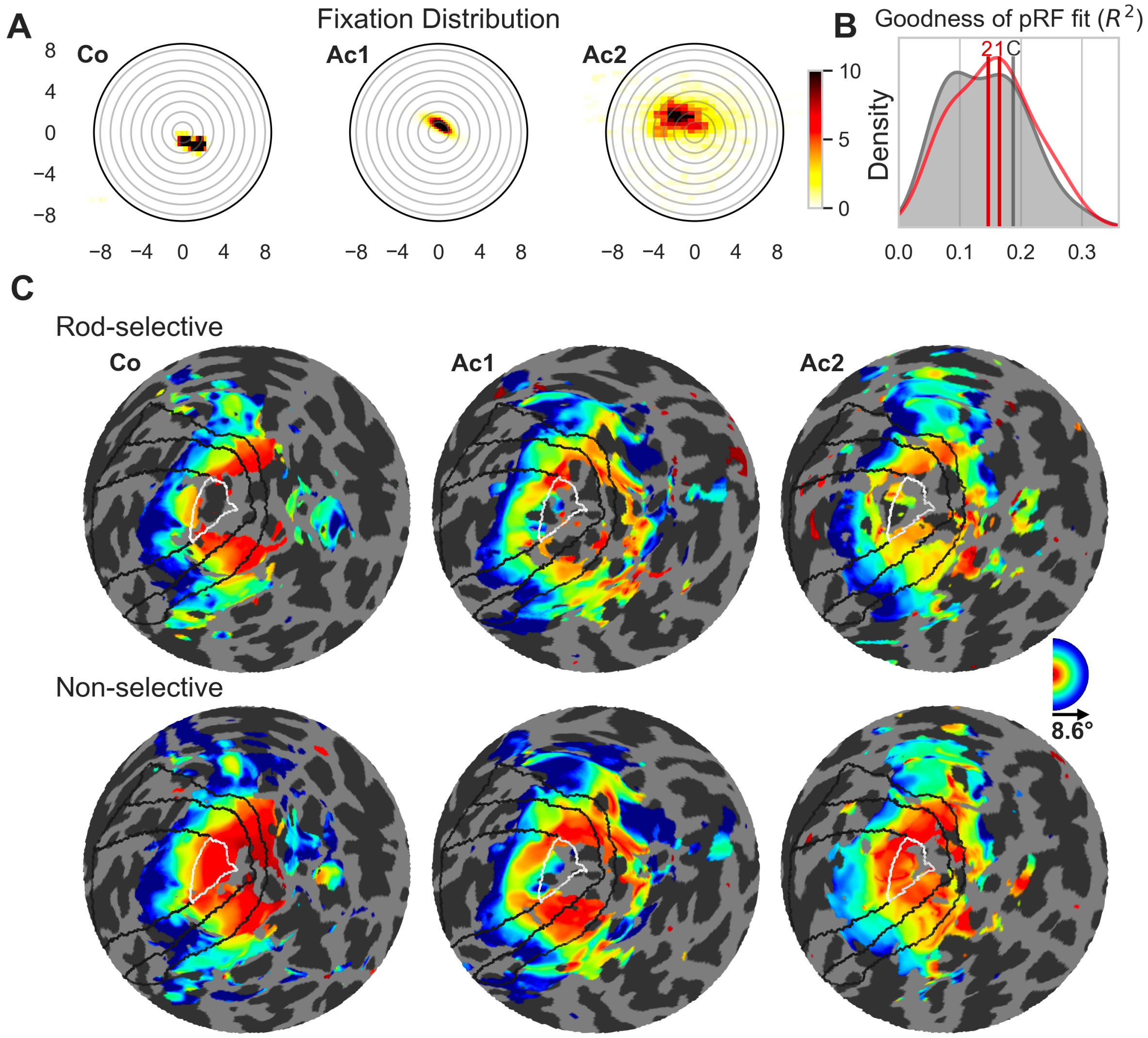


*Figure S2b. Example of 3 participants' eye data and eccentricity maps (A) Eye gaze heatmaps in visual space. External circle represents the largest eccentricities presented (8.6°), then each ring represents 1°. Heatmap units are in seconds, i.e., the darkest patches represent location where gaze was recorded for more than 10 seconds. (B) pRF model goodness of fit. Mean R2 across the entire foveal confluence (without filtering) was extracted for each participant. The group distribution of the means is plotted in gray for controls and red for achromats. The values of the three example participants are marked with vertical lines. C=Co, 1=Ac1, 2=Ac2. (C) Flat surface right hemisphere eccentricity maps of both stimulation conditions in individual's native space. Foveal confluence is marked with a white line.*

Figure S2b shows examples for how greater eye movement does not directly correspond to reduced foveal coverage. "Co" in the figure is a control with equivalent fixation stability to that of Ac1, one of the best fixators in the achromat group (Figure S2b panel A). Even with a fairly good fixation in Ac1 we do not see a filling in to the same extent as Co (Figure S2b panel C). Ac2 however, had a relatively poorer fixation (Figure S2b panel A), however they had the largest coverage (Figure S2b panel C). Each of these individuals had relatively comparable $R^2$ values (Figure S2b panel B).

#### Supplementary Material 3. Modelling the rod scotoma

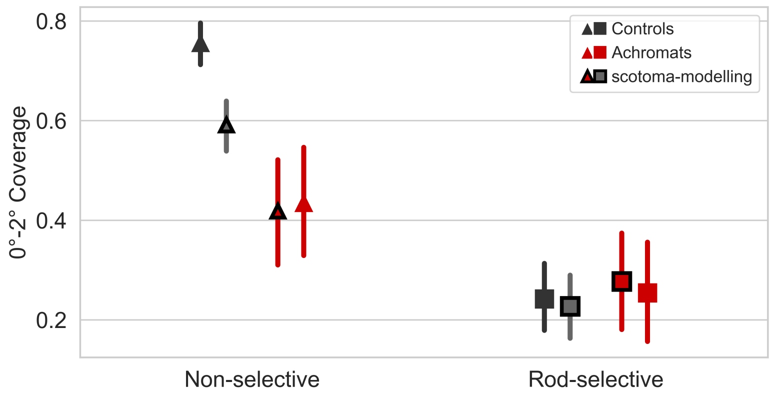


*Figure S3A. Foveal projection zone (0°-2°) coverage with and without modelling the rod-scotoma. Coverage is the fraction of vertices within the foveal confluence that contain visual information (i.e., pRF $R^2$>0.1). Triangle markers: group means for the non-selective condition, square markers: group means for the rods only condition. Black markers: Controls group means, red markers: achromat group means. Black/red vertical lines around the mean: 95% confidence interval.*

Studies on artificial scotomas, where part of the visual field is masked, suggest that pRF estimates of eccentricity and size can be biased by fitting scotoma-edge artefacts, and that these can be partially mitigated by modelling the scotoma in the pRF fitting procedure (e.g., Binda et al. 2013).

We therefore repeated the pRF modelling procedure with the rod-scotoma being modelled as a black oval mask (1.25°x0.96°) over the stimulus aperture model. As expected, a visible difference between the two models is only apparent in the non-selective condition in controls where the cones in the rod-free zone are being stimulated. In all the other conditions (rod-selective in controls, and both stimulation conditions in achromats) only the rods are stimulated, therefore the masked stimulus still matches the retinal activation, and no major differences can be observed. Performing the same statistical tests applied to the full model in the main text yields equivalent results of equivalent coverage in the rod-selective condition, with equivalent coverage across groups($t_{(47)}$ = *0.78, p=0.43,* $BF_{10}$*=0.31*) and controls show a higher coverage in the non-selective stimulation condition compared to achromats (Mann $U_{(52)}$*=141, p<0.01; unequal variance, reverted to non-parametric*).

This consistency in pRF properties when modelling the rod scotoma, is in line with previous results from scotoma modelling; While Binda and colleagues found that this normalised pRF shifts, others found no effect (Haak et al. 2012, Prabhakaran et al. 2020). Notably, the rod-free zone in achromatopsia is considerably smaller (~0.5° radius) than most tested artificial scotomas, and as artificial scotomas (screen-based masking) are not equivalent to retinal scotomas from inactive photoreceptors, it is unclear how artificial scotoma findings generalise to clinical populations. Our results are in line with a recent achromatopsia study (Anderson et al. 2024) which also found no change in pRF estimates with scotoma modelling.

Nonetheless, we cannot completely rule out scotoma-edge artefacts, as these are notoriously hard to account for.

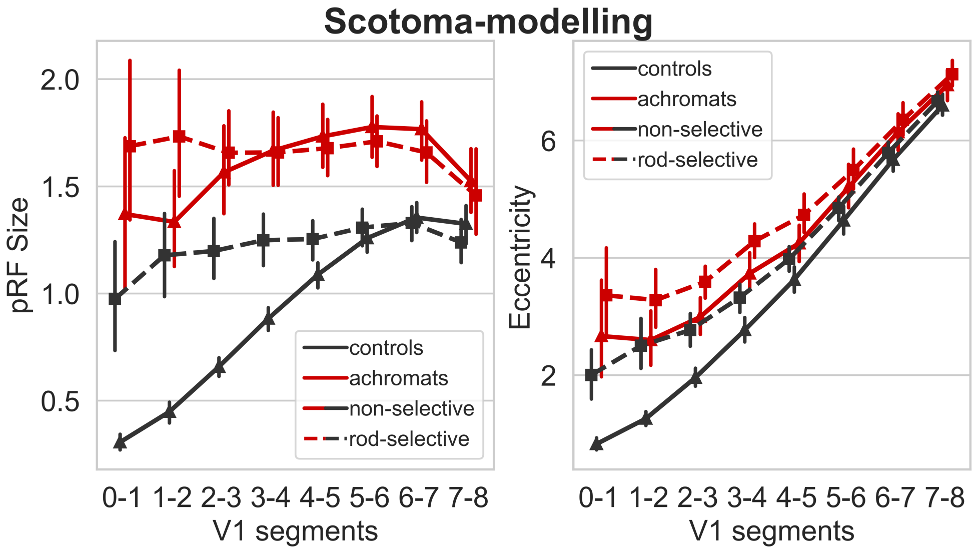


*Figure S3B. pRF sizes and eccentricity along the atlas-defined V1 segments when modelling the rod-scotoma. Eccentricity and pRF size effects remain consistent. Black/red vertical lines around the mean represent the 95% confidence interval.*

Differences between the two analyses are minute and produce equivalent effects to those described in the main text:

- pRF sizes are larger in achromats compared to controls across all non-foveal V1 segments in both conditions.

  Rod-selective condition:

  2°-3°: $t_{(48)}=3.59$, $p_{unc}=0.001$; 3°-4°: $t_{(48)}=3.50$, $p_{unc}=0.001$; 4°-5°: $t_{(48)}=4.66$, $p_{unc} <0.001$; 5°-6°: $t_{(48)}=4.80$, $p_{unc}<0.001$; 6°-7°: $t_{(48)}=4.00$, $p_{unc}<0.001$; 7°-8°: $t_{(48)}=2.06$, $p_{unc}=0.045$

  Non-selective condition:

  2°-3°: $t_{(52)}=12.11$, $p_{unc}<0.001$; 3°-4°: $t_{(52)}=10.92$, $p_{unc}<0.001$; 4°-5°: $t_{(52)}=9.31$, $p_{unc}<0.001$; 5-6: $t_{(52)}=7.34$, $p_{unc}<0.001$; 6-7: $t_{(52)}=5.59$, $p_{unc}<0.001$; 7-8: $t_{(52)}=2.23$, $p_{unc}=0.030$

- Eccentricities are slightly higher in achromats compared to controls in the rod-selective condition:

  2°-3°: $t_{(48)}=3.13$, $p_{unc}=0.003$; 3°-4°: $t_{(48)}=4.25$, $p_{unc}<0.001$; 4°-5°: $t_{(48)}=3.55$, $p_{unc}=0.001$; 5°-6°: $t_{(48)}=3.18$, $p_{unc}=0.003$; 6°-7°: $t_{(48)}=3.21$, $p_{unc}=0.002$; 7°-8°: $t_{(48)}=2.77$, $p_{unc}=0.008$;

#### Supplementary Material 4. Connective field analysis model fit

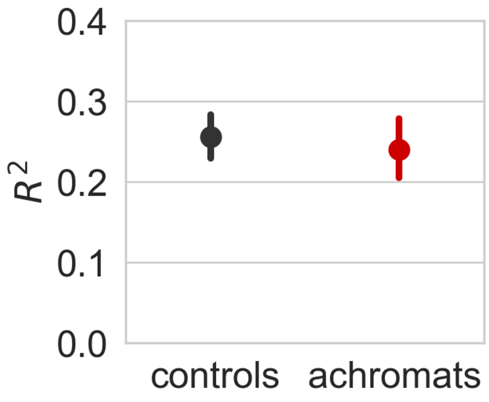


*Figure S4. Goodness of CF fit across all (unfiltered) V3 vertices (CF R2). Dots represent the group mean, Black/red vertical lines around the mean represent the 95% confidence interval.*

Goodness of CF fit was not significantly different across the two groups ($t_{(47)}$=0.63, p=0.53, $BF_{10}$=0.36) for the rod-selective condition.

#### Supplementary Material 5. pRF mapping stimuli

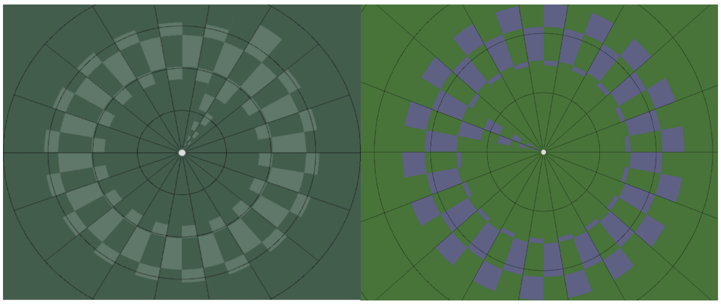

*Figure S5. Example of non-selective stimuli (left), and rod-selective stimuli (right) used in pRF mapping paradigm.*

#### Supplementary Material 6. pRF results under larger $R^2$ thresholds

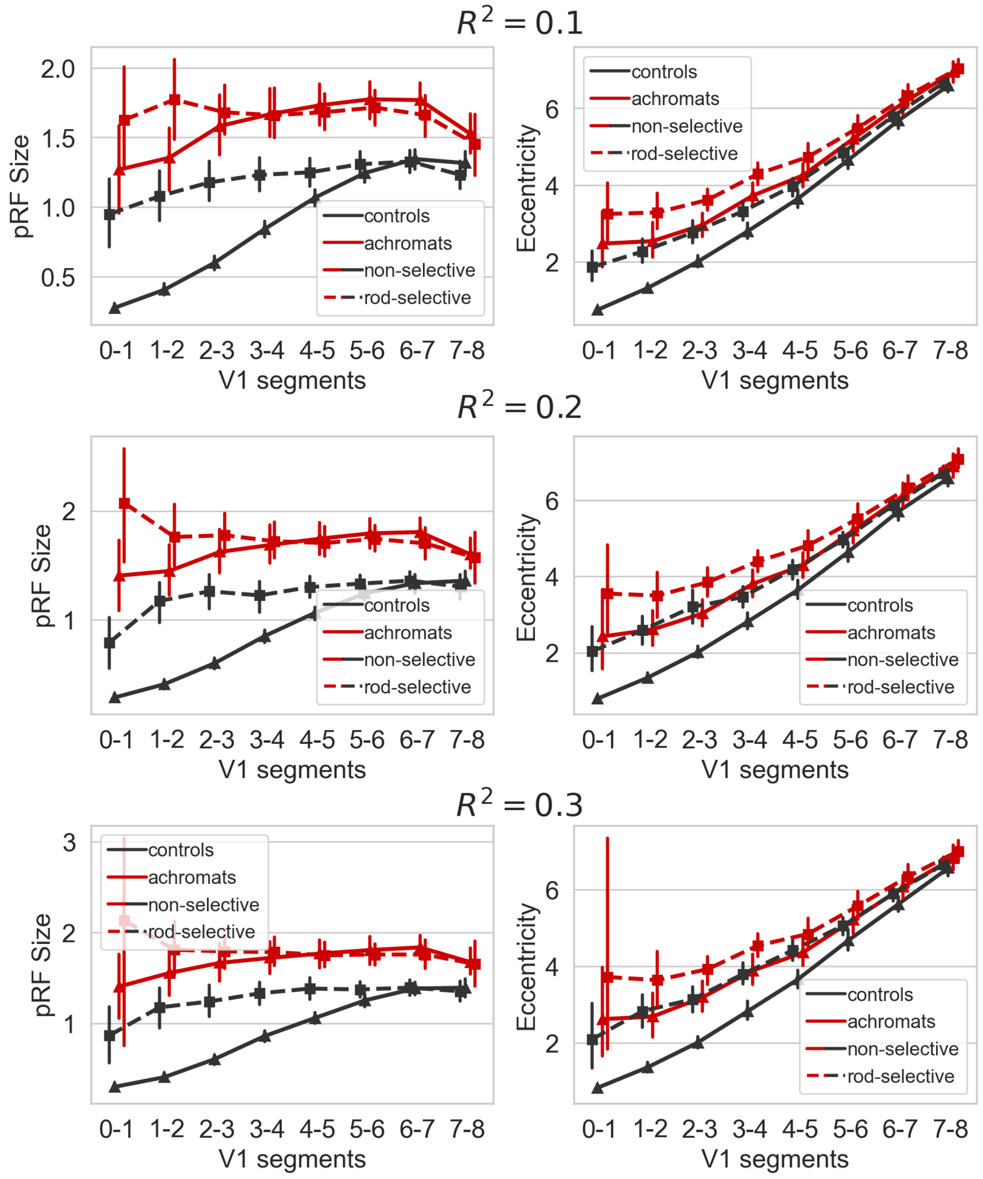


*Figure S6. pRF size and eccentricity results under various $R^2$ thresholds showing identical patterns.*

#### Supplementary Material 7. V3 and V1 pRF size results by CF centre

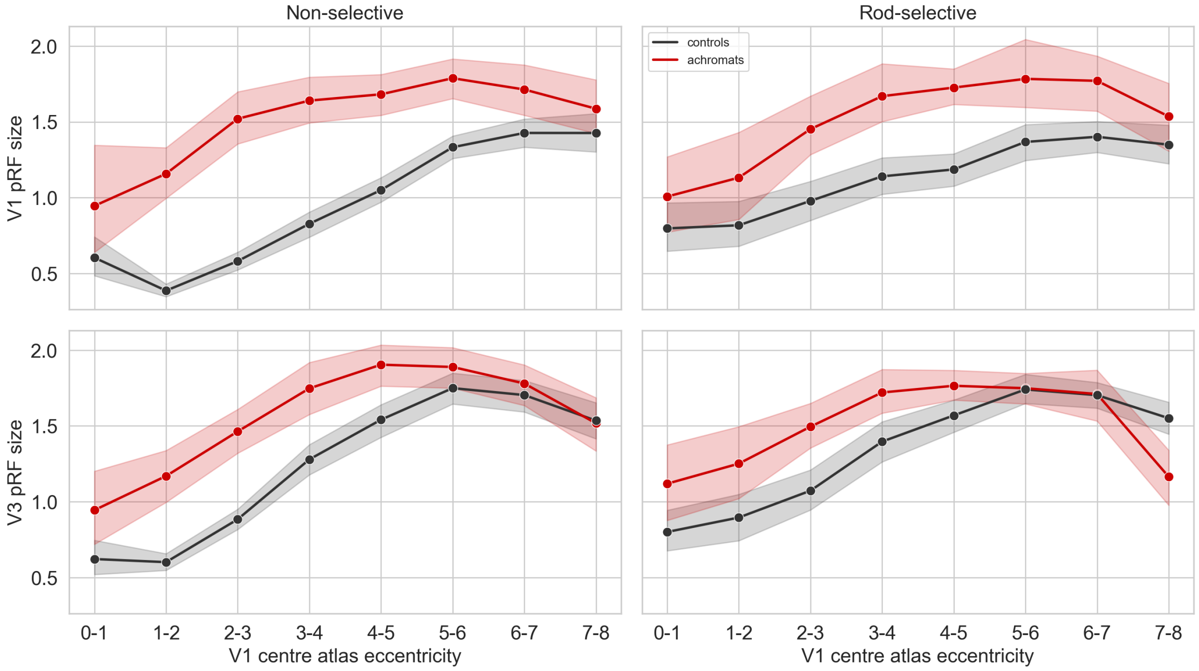


*Figure S7. pRF size results for visual area V3 and their V1 centres. For each vertex in V3, the pRF size of its V1 CF centre (top row), and its own pRF size (bottom row). Results are then binned based on the V1 centre atlas eccentricity and averaged across participants and groups. Shaded area: 95% confidence interval.*
